# BLTP1-dependent phospholipid efflux prevents mitochondrial lipid overload and apoptosis at FKBP8-tethered ER-mitochondrial contact

**DOI:** 10.1101/2025.09.30.679455

**Authors:** Qingzhu Chu, Xinran Lu, Tiansi Cui, Zhaomin Zhou, Jun Luo, Weiping Chang, Lin Deng, Wei-Ke Ji

## Abstract

Maintenance of mitochondrial phospholipid homeostasis is critical for organellar function and cell survival, yet mechanisms enabling phospholipid efflux from mitochondria remain undefined. We identify BLTP1 as a phospholipid transporter that exports phospholipids from mitochondria to maintain lipid homeostasis. The outer mitochondrial membrane protein FKBP8 recruits BLTP1 to ER-mitochondrial contact sites, establishing a dedicated lipid export pathway. BLTP1 deficiency triggers pathological accumulation of phosphatidic acid (PA), phosphatidylglycerol (PG), and cardiolipin (CL) within mitochondria. This lipid overload is associated with elevation of mitochondrial reactive oxygen species (ROS), bioenergetic dysfunction, and apoptosis. Critically, depleting intramitochondrial lipid transfer proteins (e.g., PRELID1) or enzymes in the CL synthesis pathway (e.g., PTPMT1, CRLS1) prevents apoptosis caused by BLTP1 deficiency. Our findings defines a BLTP1-FKBP8-dependent mechanism for mitochondrial phospholipid efflux that is essential for mitochondrial function and cell survival.

## Introduction

Phospholipids constitute the fundamental structural components of cellular membranes, essential for maintaining their integrity, fluidity, and dynamic barrier functions. Mitochondria, central hubs for cellular energy metabolism and homeostasis, critically depend on tightly regulated phospholipid metabolism to maintain their structural and functional integrity. This regulation encompasses phospholipid synthesis, degradation, remodeling, and inter-organellar trafficking^1^. Disruption of mitochondrial phospholipid homeostasis impairs core organelle functions, adversely impacting diverse processes including cellular bioenergetics, metabolism, and survival^2^.

While mitochondria primarily acquire phospholipids via lipid transfer proteins (LTPs)- mediated transfer from the endoplasmic reticulum (ER), they also synthesize some phospholipids internally. Key mitochondrial phospholipids include phosphatidic acid (PA), phosphatidylcholine (PC), phosphatidylglycerol (PG), cardiolipin (CL), phosphatidylserine (PS), and phosphatidylethanolamine (PE). PA, a central biosynthetic precursor, originates from de novo synthesis within mitochondria or is imported from the ER to the outer mitochondrial membrane (OMM). Within the inner mitochondrial membrane (IMM), PA is converted sequentially to CDP-diacylglycerol (CDP-DAG) by CDS1/2, then to phosphatidylglycerol phosphate (PGP) by TAMM41/PGS1, and finally to PG by phosphatase PTPMT1^3, 4, 5^. PG serves as the substrate for CL synthesis catalyzed by cardiolipin synthase (CRLS1), followed by CL remodeling to its mature form^6, 7^.

Inter-organelle phospholipid transfer, essential for lipid homeostasis and organelle dynamics, occurs at membrane contact sites (MCSs) facilitated by LTPs ^8–12^ ^13–15^ ^16^ ^17, 18^. A distinct class of LTPs, termed bridge-like lipid transfer proteins (BLTPs), possess a unique architecture: a repeating β-groove (RBG) rod-like structure forming a continuous hydrophobic channel^19, 20^. BLTP1 (KIAA1109), a poorly characterized mammalian BLTP, shares homology with yeast Csf1, which localizes to ER-associated MCSs and transports PE from mitochondria to the ER for glycosylphosphatidylinositol (GPI) anchor biosynthesis^15^. BLTP1 Homologs in Drosophila (Tweek) and C. elegans (LPD-3) are implicated in phosphoinositide metabolism, synaptic vesicle endocytosis, micropinocytosis, cold tolerance, and insulin-mTOR signaling during aging^21–23^. The cellular localization and function of BLTP1 in mammals remains elusive.

Critically, loss-of-function mutations in human BLTP1 cause Alkuraya-Kucinskas syndrome, a severe neurodevelopmental disorder characterized by global developmental delay, profound brain malformations, joint deformities, and high neonatal mortality; survivors often experience significant intellectual disability and epilepsy^24^. Despite this clear genetic link, the specific cellular functions of BLTP1 in mammals and the pathogenic mechanisms underlying Alkuraya-Kucinskas syndrome remain unknown, hindering therapeutic development.

Therefore, this study aimed to define the cellular role and molecular mechanisms of BLTP1 in mammals. We report that the outer mitochondrial membrane (OMM) protein FKBP8 acts as an adaptor recruiting BLTP1 to ER-mitochondria MCSs. Deficiency of BLTP1 results in aberrant mitochondrial accumulation of phospholipids central to the CL synthesis pathway—specifically PA, PG, and CL—associating with mitochondrial reactive oxygen species (mtROS) overproduction, bioenergetic dysfunction and apoptosis. Crucially, genetic silencing of intramitochondrial lipid transfer proteins (e.g., PRELID1) or enzymes within the CL synthesis pathway (e.g., PTPMT1, CRLS1) rescues apoptosis induced by BLTP1 loss. Our findings unveil a key role of BLTP1 in phospholipid efflux from mitochondria and defines a BLTP1-dependent surveillance mechanism at FKBP8-tethered ER-mitochondria MCS for mitochondrial function and cell survival.

## Results

### FKBP8 is a novel BLTP1-interacting protein that binds to C-terminal of BLTP1

BLTP1 is an integral ER protein anchoring ER membrane via a transmembrane domain (TM) at the N-terminus (Fig. 1A & S1A). The C. elegans BLTP1 homolog LPD-3 interacts with ER-resident proteins (e.g., Spigot) via a region at its N-terminal that may facilitate lipid extraction^25^. However, the mechanism governing the association of BLTP1 with the other organelle at MCS in mammals remained unknown. To identify interactors determining the subcellular targeting of BLTP1, we generated a U2OS cell line expressing endogenously 3xFlag-tagged BLTP1 (BLTP1^Flag) using CRISPR-Cas9 (Fig. 1B & S1B). The BLTP1^Flag Knockin (BLTP1^Flag KI) line was validated by genomic PCR and small interfering RNAs (siRNA)-mediated depletion in dot blot (Fig. S1C, D).

**Fig. 1.**
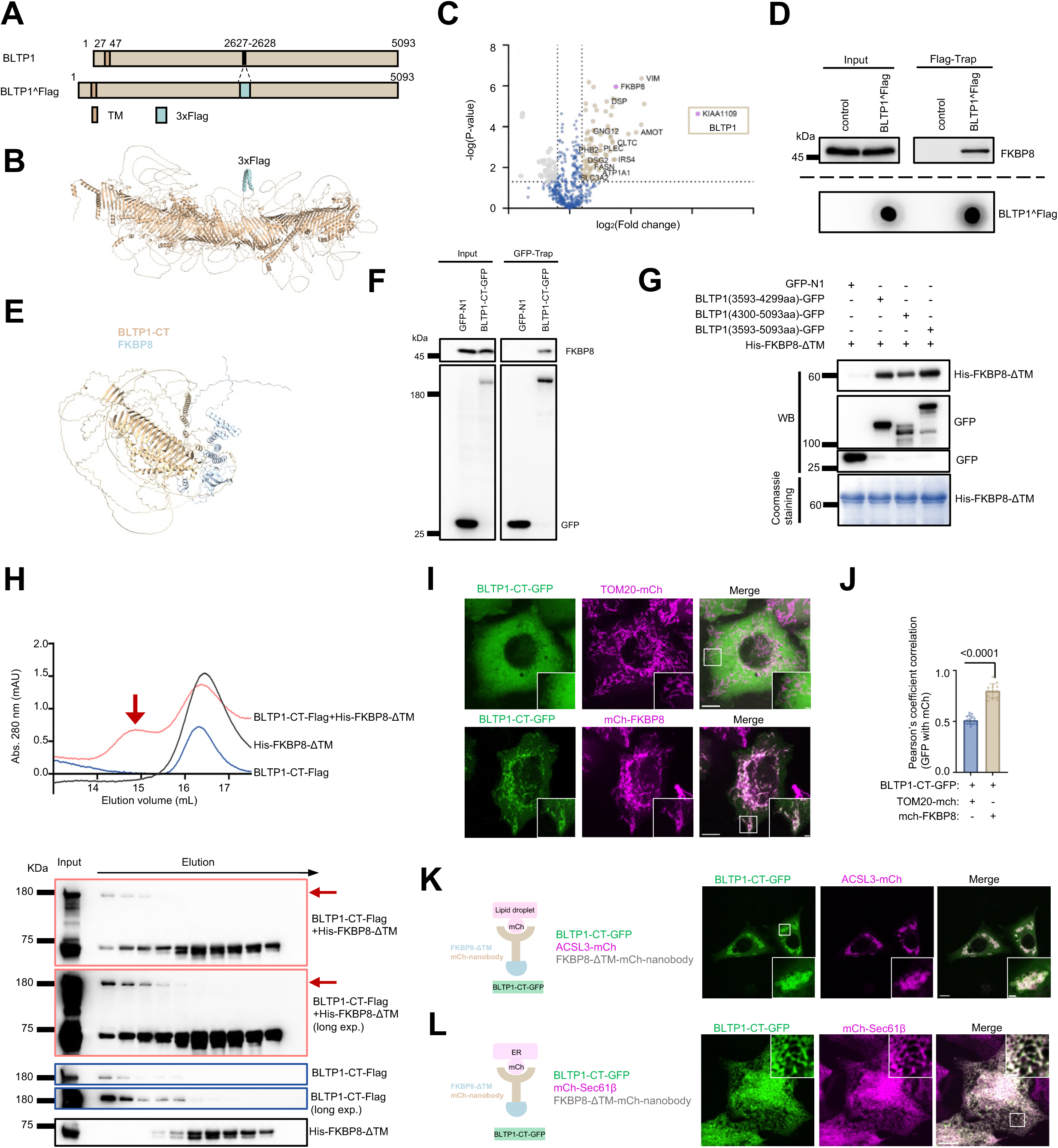
Identification of FKBP8 as a novel BLTP1 interacting protein. (A) Schematic cartoon of internally Flag tagging in the BLTP1^Flag Knock-in cell line. (B) AlphaFold-predicted structures of BLTP1^Flag with the 3x Flag tag labeled in green. (C) Volcano plot of protein candidates coIPed with BLTP1 in Knock-in cell line compared with protein candidates coIPed with vector only. Candidates that were considered significant (−log [P value] > 1.3; P < 0.05) were labeled in light brown (Log2 [fold change] > 1; increased in abundance) or gray (Log2 [fold change] < -1 decreased in abundance). (D) coIP assay showed an interaction between endogenous BLTP1 and endogenous FKBP8 in the BLTP1 Knock-in cell line. Dot blots were performed for BLTP1^Flag. In this assay, WT U2OS cells were used for the control. (E) AlphaFold Multimer prediction showing that FKBP8 binds the C-terminal region of BLTP1 (residues 3593–5093; BLTP1-CT). (F) GFP-Trap assays demonstrate an interaction between BLTP1-CT-GFP and endogenous FKBP8 in HEK293 cells. (G) In vitro pulldown assays using BLTP1-CT-GFP bound on GFP-Trap beads and purified His-FKBP8-DTM. (H) Size exclusion chromatography elution profile of BLTP1-CT-Flag (blue), His-FKBP8-ΔTM (black), and a mixture of BLTP1-CT and FKBP8-ΔTM (red). From top to bottom: immunoblots of the corresponding peak fractions from the BLTP1-CT – FKBP8-ΔTM, BLTP1-CT, and FKBP8-ΔTM elution. (I) Representative confocal images of an U2OS cell expressing BLTP1-CT-GFP (green) with Tom20-mCh (magenta; top panel) or mCh-FKBP8 (magenta; bottom panel) with insets. (J) Pearson’s correlation coefficient in cells as shown in (I). In the quantification, more than 14 cells for each condition were analyzed from three independent experiments. Two-tailed unpaired Student’s t test. Mean ± SD. (K) Left: schematic cartoon of mCh-nanobody-mediated recruitment of FKBP8-ΔTM-mCh-nb to lipid droplet membranes (ACSL3-mCh) in the presence of BLTP1-CT-GFP. Right: representative images of a U2OS cell expressing BLTP1-CT-GFP (green), ACSL3-mCh (magenta) in presence of FKBP8-ΔTM-mCh-nb with insets. (L) Left: schematic cartoon of mCh-nanobody-mediated recruitment of FKBP8-ΔTM-mCh-nb to ER membranes (mCh-sec61β) in the presence of BLTP1-CT-GFP. Right: representative images of a U2OS cell expressing BLTP1-CT-GFP (green), mCh-sec61β (magenta) in presence of FKBP8-ΔTM-mCh-nb with insets. Scale bar, 10 μm in the whole cell images and 2 μm in the insets in (I, J, K).

Co-immunoprecipitation (co-IP) of BLTP1^Flag followed by mass spectrometry identified FKBP8—an outer mitochondrial membrane (OMM) protein—as one of top candidates (Fig. 1C). Endogenous co-IP confirmed a robust BLTP1-FKBP8 interaction (Fig. 1D).

Contrary to BLTP1, other BLTP proteins (BLTP2 and VPS13D) failed to be co-pelleted with FKBP8 (Fig. S1E). Consistently, BLTP2 tagged with green fluorescent protein (GFP) or Halo tagged VPS13A did not substantially co-localize with mCherry-tagged FKBP8 (mCh-FKBP8) (Fig. S1F). These results suggest a specific interaction between BLTP1 and FKBP8.

We then sought to determine which region of BLTP1 is responsible for interacting with FKBP8. To begin with, we employed AlphaFold Multimer to predict that FKBP8 binds the C-terminal region of BLTP1 (residues 3593–5093; BLTP1-CT) (Fig. 1E), which was further confirmed by GFP Trap assays (Fig. 1F). Notably, GFP-Trap assays further showed that BLTP1 preferentially interacted with FKBP8 via the CT region (Fig. S1G).

Next, we asked whether BLTP1 directly interacts with FKBP8 using *in vitro* pull-down assay. As full-length BLTP1 proved refractory to purification, we used the purified BLTP1-CT fragment in the assay. After removing potential binding partners by rigorous washing using high-salt (500 mM NaCl) buffer, the BLTP1-CT fragment was incubated with purified FKBP8 with a deletion of TM domain (His-FKBP8-1′TM). Importantly, BLTP1-CT-GFP, but not GFP alone, bound to purified His-FKBP8-1′TM (Fig. 1G).

To investigate whether BLTP1 forms a stable complex with FKBP8, we employed size-exclusion chromatography (SEC). In this assay, we observed that the purified BLTP1-CT-Flag was co-eluted with His-FKBP8-1′TM (Fig. 1H), indicating of forming a stable BLTP1-FKBP8 complex in vitro.

Importantly, the BLTP1-CT was diffused in the cytosol without membrane association in U2OS cells expressing TOM20-mCh (Fig. 1I; top panel); in contrast, expression of mCh-FKBP8 evidently recruited the BLTP1 fragment to the OMM (Fig. 1I bottom panel; 1J). These results demonstrated that the CT region of BLTP1 is responsible for interacting with FKBP8.

On the other hand, we investigated how FKBP8 interacted with BLTP1. FKBP8 contains a disordered region at the NT, followed by a PPIase domain, a TPR domain and a TM domain at the CT^26^ (Fig. S2A). GFP-Trap assays indicated that deletion of the NT region or the TPR domain, but not the PPIase domain, substantially reduced the interaction with the BLTP1-CT (Fig. S2B), which was further confirmed by imaging results using co-localization analyses (Fig. S2C, D). Notably, neither the FKBP8-NT (mCh-FKBP8-NT-TM) nor the TPR domain (mCh-FKBP8-TPR-TM) alone was able to recruit the BLTP1-CT (Fig. S2C, D, bottom panels), suggesting that both of the two domains of FKBP8 are required for interaction with BLTP1.

Taking advantage of a specific interaction between mCh and an mCh nanobody (mCh-nb), we performed knock-sideways assays to definitively confirm the binding between BLTP1-CT and FKBP8. Redirecting FKBP8-ΔTM-mCh-nb to lipid droplets (via ACSL3-mCh; Fig. 1K) or ER membranes (via mCh-Sec61β; Fig. 1L) efficiently recruited the cytosolic BLTP1-CT to these membrane compartments. Together, we conclude that FKBP8 is a new BLTP1 interacting protein that binds to the CT of BLTP1.

### FKBP8 recruits BLTP1 to ER-mitochondria contact sites

Given that BLTP1 is a potential lipid transfer protein functioning at MCS, we tested whether the OMM protein FKBP8 is able to recruit BLTP1 to ER-mitochondrial MCS. Immunofluorescence and membrane fractionation in U2OS cells confirmed endogenous FKBP8 enrichment in mitochondrial-associated membranes (MAM; equivalent to ER-mitochondrial MCSs) (Fig. 2A, B), consistent with previous studies^27^.

**Fig. 2.**
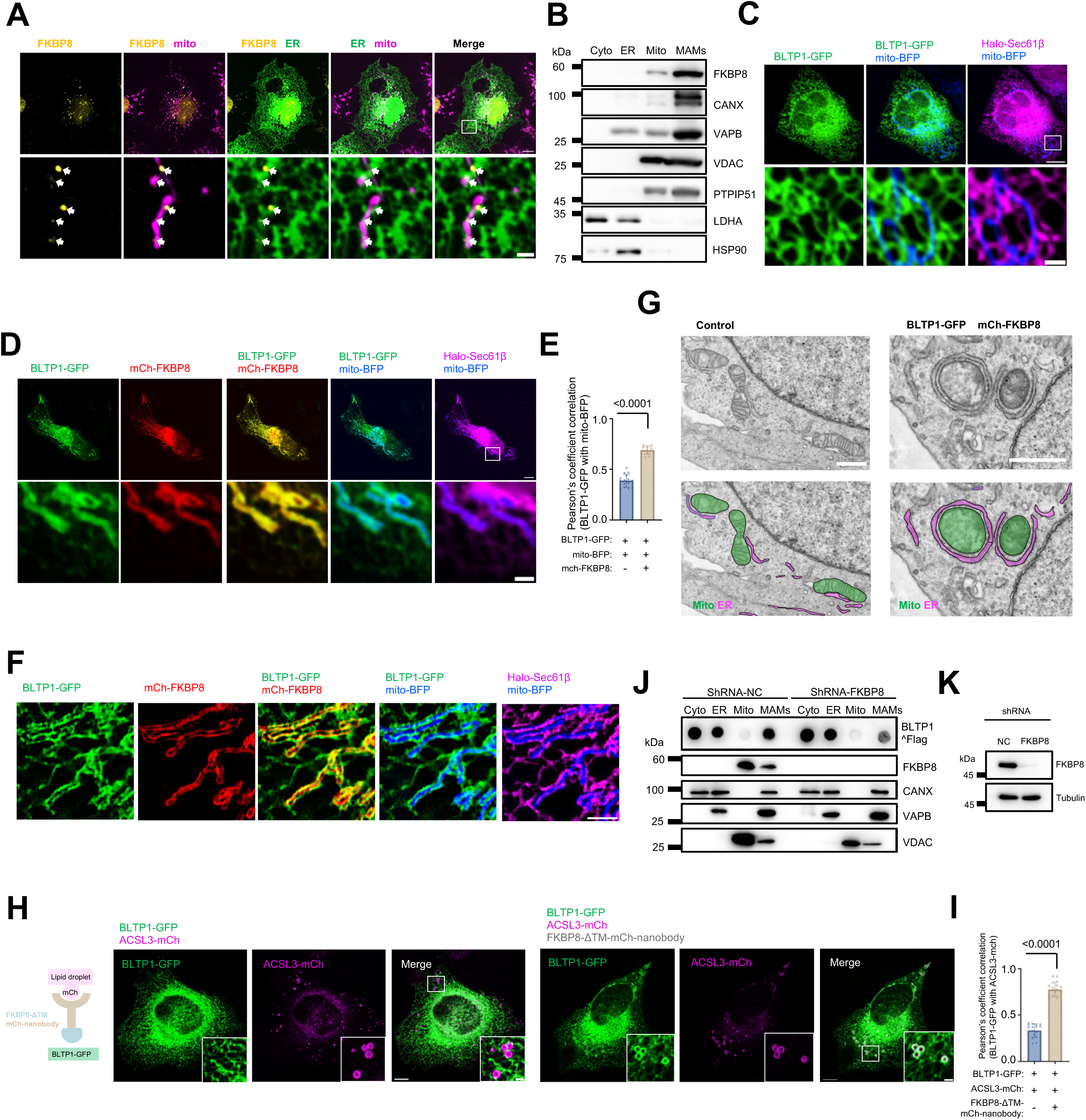
FKBP8 recruits BLTP1 to ER-mitochondrial MCS. (A) Representative images of a fixed U2OS cell stained with FKBP8 antibody (yellow), the ER (green) and mitochondria (magenta) with an inset at cell periphery. Arrows denote FKBP8 puncta at ER-mitochondrial junctions. (B) Representative western blots showing the enrichment of endogenous FKBP8 in the MAM membrane fraction isolated from U2OS cell lysates. (C, D) Confocal images show an U2OS cell expressing BLTP1-GFP (green), Halo-Sec61β (magenta), and mito-BFP (blue) in absence (C) or presence (D) of mCh-FKBP8(red) with insets. (E) Pearson’s correlation coefficient in cells as shown in (C, D). In the quantification, more than 10 cells for each condition were analyzed from three independent experiments. Two-tailed unpaired Student’s t test. Mean ± SD. (F) Representative super-resolution SIM micrograph of an U2OS cell expressing BLTP1-GFP (green), mCh-FKBP8 (red), Halo-Sec61β (magenta), and mito-BFP (blue). (G) Representative TEM micrographs showing examples of ER-mitochondrial MCSs in vector (left panel) or BLTP1-GFP and mch-FKBP8 co-expressing U2OS cells (right panel). Mitochondria and the ER were labeled with green or magenta, respectively. (H) Left: schematic cartoon of mCh-nb-mediated recruitment of FKBP8-ΔTM-mCh-nb to lipid droplet membranes (ACSL3-mCh) in presence of BLTP1-GFP. Right: representative images of U2OS cells expressing BLTP1-GFP (green) and ACSL3-mCh (magenta) in absence or presence of FKBP8-ΔTM-mCh-nb with insets. (I) Pearson’s correlation coefficient in cells as shown in (H). In the quantification, more than 15 cells for each condition were analyzed from three independent experiments. Two-tailed unpaired Student’s t test. Mean ± SD. (J) Representative western blots showing the distribution of endogenous BLTP1^Flag in membrane fractions isolated from BLTP1 Knock-in cell line cell lysates under control or FKBP8 depletion. (K) Representative western blots confirming shRNA-mediated FKBP8 depletion in cells as shown in (J). Scale bar, 10 μm in the whole cell images and 2 μm in the insets in (A, C, D, H), 2 μm in (F) and 1 μm in (G).

Live-cell imaging confirmed that exogenous BLTP1-GFP was uniformly distributed over the ER network without evident enrichment at junctions between the ER and mitochondria (Fig. 2C). Strikingly, mCh-FKBP8 overexpression recruited BLTP1-GFP to sites of extensive ER-mitochondria apposition (Fig. 2D, E), likely representing dramatically elevated ER-mitochondrial MCSs.

To validate the result, we performed super-resolution structured illumination microscopy (SIM). The SIM micrographs clearly showed that BLTP1-GFP was strongly colocalized with mCh-FKBP8 at sites where the ER tightly wrapped the OMM (Fig. 2F). These results confirmed that FKBP8-mediated BLTP1 recruitment coincided with expanded ER-mitochondria MCS.

Consistent with these light microscopy results, transmission electron microscopy (TEM) demonstrated a strong increase in ER-mitochondria contacts in cells co-expressing BLTP1-GFP and mCherry-FKBP8 relating to cells expressing empty vectors (Fig. 2G).

In addition, we investigated whether FKBP8 is able to recruit BLTP1 at endogenous level in the BLTP1^Flag KI line. We noticed that the endogenous BLTP1^Flag formed dim puncta and the specificity of BLTP1^Flag was validated by siRNA-mediated depletion using anti-Flag antibody in immunofluorescence staining (Fig. S3A, B). The cellular localization of endogenous BLTP1 relative to the ER was then examined (Fig. S3C, D); contrary to overexpressed BLTP1, endogenous BLTP1^Flag formed puncta on the ER (Fig. S3C). Notably, approximate 20% of these BLTP1 puncta was located to ER-mitochondrial MCSs (Fig. S3E, F), consistent with yeast Csf1 localization^15^. Importantly, upon expression of mCh-FKBP8, over 60% of endogenous BLTP1^Flag puncta were associated with junctions between the ER and mitochondria (Fig. S3G, H), indicating that FKBP8 promotes the recruitment of endogenous BLTP1 to ER-mitochondrial MCS.

Furthermore, we investigated whether the BLTP1-FKBP8 interaction is sufficient to mediate the formation of inter-organelle MCS by the knock-sideway assays (Fig. 2H, left panel), as described above. The colocalization between BLTP1 and lipid droplets was neglectable in the control (Fig. 2H, middle panel). In contrast, BLTP1-GFP was dramatically recruited to lipid droplets in presence of FKBP8-ΔTM-mCh-nb, inducing extensive ER-LD contacts (Fig. 2H, right panel; 2I).

Next, we asked whether FKBP8 is required for the localization of BLTP1 to ER-mitochondrial MCS. Membrane fractionation revealed a significant portion of endogenous BLTP1^Flag in the MAM fraction (Fig. 2J), consistent with our imaging results. Notably, it was significantly reduced upon small hairpin RNA (shRNA)-mediated FKBP8 depletion (Fig. 2J, K). Notably, the level of VAPB was not substantially impacted, ruling out the possibility that the reduction of BLTP1 in MAMs is due to impaired ER-mitochondrial interactions caused by FKBP8 deficiency. Therefore, we conclude that the interaction with FKBP8 is sufficient and required for BLTP1 localization at ER–mitochondrial MCSs.

### BLTP1 depletion disrupts mitochondrial dynamics and function

Since our results demonstrated BLTP1 as an ER-mitochondrial MCS protein, we then sought to explore the cellular function of BLTP1 at these sites. Depletion of either BLTP1 or FKBP8 mediated by siRNAs resulted in mitochondrial fragmentation (Fig. 3A, B, E, F). The observed effect of BLTP1 on mitochondrial dynamics is specific, as demonstrated by the consistent phenotype resulting from two independent siRNAs (#1 and #3). We thus utilized siRNA #3 for BLTP1 depletion in all experiments unless stated otherwise. To further validate these results, we performed TEM. TEM micrographs showed that mitochondria were significantly shortened in BLTP1-depleted cells (Fig. 3C, D).

**Fig. 3.**
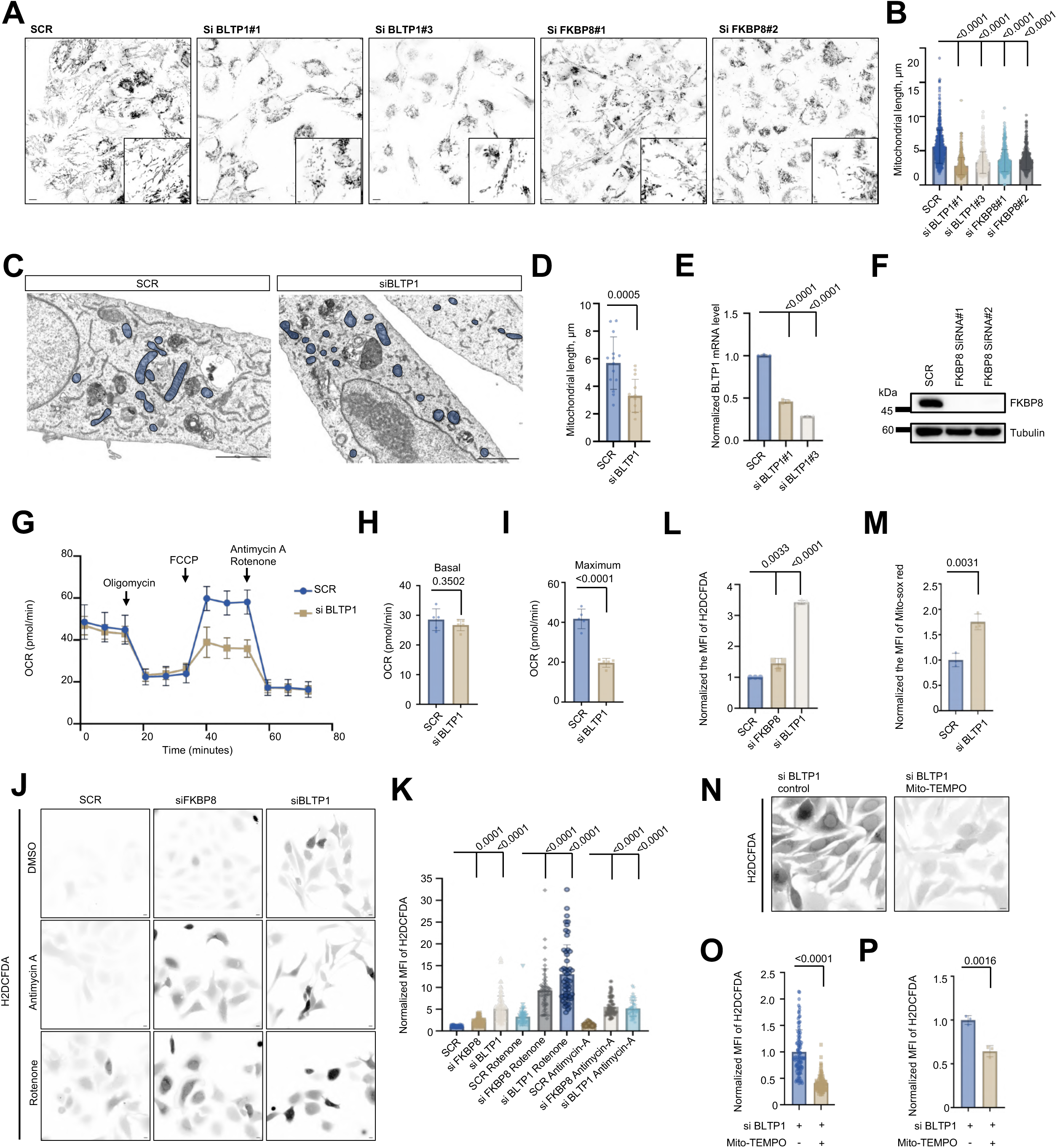
BLTP1 depletion disrupts mitochondrial dynamics and function. (A) Representative live-cell confocal images of control, BLTP1 or FKBP8 depleted U2OS cells stained with Mito-tracker with insets. (B) Quantification of mitochondrial length in cells as in (A). More than 34 cells for each condition were analyzed from three independent experiments. Two-tailed unpaired Student’s t test. Mean ± SD. (C) Representative TEM micrographs showing examples of mitochondria in control or BLTP1 depleted U2OS cells. Mitochondria were manually labeled in blue. (D) Quantification of mitochondrial length in cells as in (C). More than 10 cells for each condition were analyzed. Two-tailed unpaired Student’s t test. Mean ± SD. (E) qPCR assays showing the efficient of siRNA-mediated suppression in BLTP1 from three independent experiments. Two-tailed unpaired Student’s t test. Mean ± SD. (F) Representative western blots demonstrating efficiency of siRNA-mediated FKBP8 depletion. (G-I) Mitochondrial OCR measured in control or BLTP1 depleted U2OS cell. OCR trace was obtained by sequential measurement of basal OCR, OCR after the addition of Oligomycin, FCCP, and Rotenone/Antimycin A from 5 independent assays. (H, I) The basal (H) or maximum OCR after FCCP addition (I) as in (G). Two-tailed unpaired Student’s t test. Mean ± SD. (J) Representative images of control, BLTP1 or FKBP8 depleted U2OS cells stained with H2DCFDA under DMSO (top), Antimycin A (middle) or Rotenone (bottom) treatment. (K) Normalized H2DCFDA fluorescence intensity of control, BLTP1 or FKBP8-depleted cells as in (J). More than 50 cells for each condition were analyzed. Two-tailed unpaired Student’s t test. Mean ± SD. (L) Flow cytometry analyses of H2DCFDA fluorescence intensity in control, BLTP1 or FKBP8-depleted U2OS cells from three independent assays. The H2DCFDA fluorescence intensity was normalized to that of control cells. More than 10000 cells for each condition were analyzed. Two-tailed unpaired Student’s t test. Mean ± SD. (M) Normalized Mito-SOX red fluorescence intensity of control or BLTP1 -depleted cells by flow from three independent assays. More than 10000 cells for each condition were analyzed. Two-tailed unpaired Student’s t test. Mean ± SD. (N) Representative images of the H2DCFDA fluorescence intensity of BLTP1-depleted U2OS cells in absence or presence of Mito-TMEPO (200 μM). (O) Normalized H2DCFDA fluorescence intensity of cells as in (N). (P) Flow cytometry analyses of the H2DCFDA fluorescence intensity of BLTP1-depleted cells in absence or presence of 200 μM Mito-TMEPO treatment from three independent assays. The H2DCFDA fluorescence intensity was normalized to that of control cells. More than 10000 cells for each condition were analyzed. Two-tailed unpaired Student’s t test. Mean ± SD. Scale bar, 10 μm in the whole cell images and 2 μm in the insets in (A) and 10 μm in (J, N). 2 μm in (C).

Mitochondrial fragmentation is a hallmark of mitochondrial stress and dysfunction^28^. We therefore investigated whether BLTP1 deficiency impairs mitochondrial function. Seahorse analyses revealed a strong reduction in maximum, but not basal, oxygen consumption rates in BLTP1-depleted cells (Fig. 3G, H, I), indicating that BLTP1 depletion results in bioenergetic dysfunction.

Importantly, both BLTP1 and FKBP8 suppression significantly elevated cellular ROS, as shown by intensity of a fluorescent ROS probe H2DCFDA (Fig. 3J, K), consistent with a reported role of LPD-3, the C. elegans homolog of BLTP1, implicated in ROS signaling^22^. Notably, the effect was exacerbated by mitochondrial poisons rotenone and Antimycin A (Fig. 3J, middle & bottom; 3K), suggesting that ROS generated in cells depleted of BLTP1 is derived from mitochondria. To validate these results, we also performed high-throughput flow cytometry assays, and confirmed that depletion of either BLTP1 or FKBP8 significantly elevated cellular ROS (Fig. 3L).

Furthermore, we generated BLTP1 knockout HeLa cells (BLTP1 KO) by CRISPR-Cas9 (Fig. S4A), and two BLTP1 KO clones (#20 & #27) were validated by genomic DNA analysis (Fig. S4B) and quantitative PCR (Fig. S4C). Consistent with siRNA-mediated depletion, a significant increase in ROS was observed in these two independent clones either by imaging (Fig. S4D, E, F & G) or flow cytometry (Fig. S4H), which was further boosted by mitochondrial stress (rotenone and antimycin A). Notably, we noticed that the efficiency of the BLTP1 KO gradually decreased after multiple rounds of cell passaging due to cell death (see next section of this manuscript). Therefore, we used siRNA-mediated acute depletion for all subsequent loss-of-function experiments in this study.

Importantly, flow cytometry showed that BLTP1 deficiency caused a significant increase in intensity of mitoSOX (Fig. 3M), a mitochondria-specific ROS sensor. Importantly, Mito-TEMPO, a mitochondria-targeted antioxidant, significantly reversed the ROS increase in BLTP1-depleted cells either by imaging (Fig. 3N, O) and flow cytometry (Fig. 3P), indicating that BLTP1 depletion caused an increase in mitochondrial ROS. Together, we conclude that BLTP1 depletion resulted in severe mitochondrial dysfunctions.

### BLTP1 depletion specifically triggers apoptosis

As mentioned earlier, we observed that a portion of BLTP1-depleted U2OS cells always died, indicating that BLTP1 depletion leads to cell death. To investigate which type of cell death is induced by BLTP1 deficiency, we examined the rate of cell death in presence of cell death inhibitors using flow cytometry-based SYTOX Green assay, a dead cell-specific dye. Indeed, SYTOX Green staining exhibited significant cell death in BLTP1-depleted cells (Fig. 4A), selectively prevented by the apoptosis inhibitor ZVAD-FMK—but not necroptosis (Nec-1s) or ferroptosis (Fer-1) inhibitors (Fig. 4B, C, D & F). These results indicate that BLTP1 depletion specifically leads to apoptosis. Interestingly, the antioxidant Mito-TEMPO partially rescue death (Fig. 4E, F), indicating that elevated mitochondrial ROS contributes to apoptosis induced by BLTP1 deficiency.

**Fig. 4.**
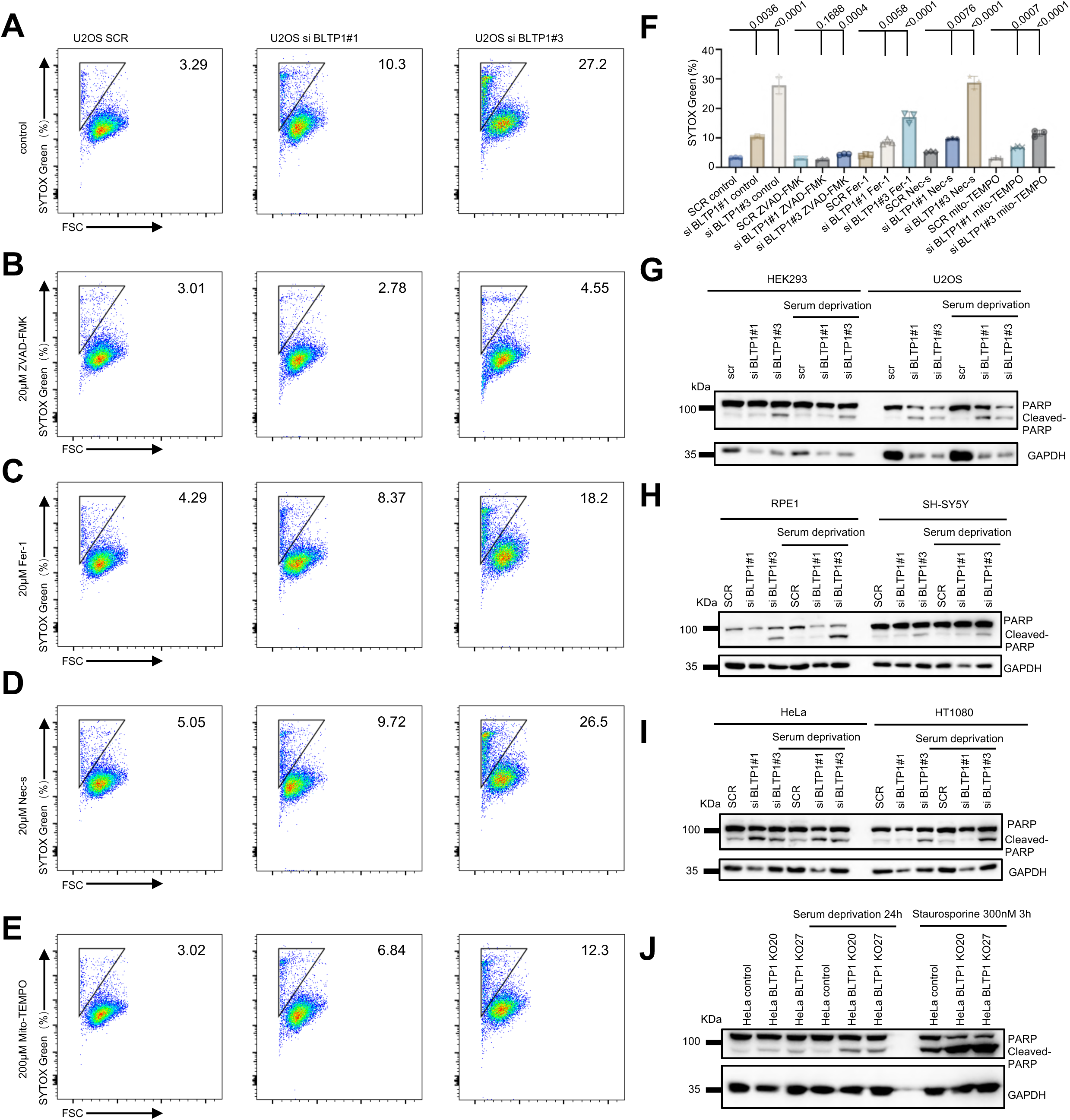
BLTP1 depletion specifically triggers apoptosis. (A) Flow cytometry analyses showing percentage of SYTOX-Green-positive U2OS cells upon scrambled or BLTP1 siRNA treatments. (B-E) As in (A), the percentage of SYTOX-Green-positive BLTP1-depleted cells under ZVAD-FMK (B), Nec-1s (C), Fer-1 (D) and Mito-TEMPO (E) treatment. (F) Quantification of percentage of SYTOX-Green-positive cells as in (A-E). More than 10000 cells for each condition were analyzed from three independent experiments. Two-tailed unpaired Student’s t test. Mean ± SD. (G) Representative western blots showing the level of PARP and Cleaved-PARP in control or BLTP1-depleted HEK293 and U2OS cells under normal or serum starvation. (H, I) Representative Western blots showing PARP/Cleaved-PARP treated with scrambled or BLTP1 siRNA in other cell lines including RPE1 and SH-SY5Y (H) or HeLa and HT1080 (I) under normal or serum starvation. (J) Representative western blots showing PARP/Cleaved-PARP in control or two BLTP1 KO HeLa clones under normal, serum starvation or staurosporine treatment.

This conclusion was further supported by Immunoblotting showing increased cleaved PARP, a hallmark of apoptosis, in serum-starved BLTP1-depleted HEK293, U2OS, RPE1, SH-SY5Y, HeLa and HT1080 cells (Fig. 4G, H, I). The two BLTP1-KO HeLa clones (KO-20, KO-27) similarly showed enhanced PARP cleavage upon serum deprivation or apoptosis inducer Staurosporine (Fig. 4J).

### BLTP1^-/-^ mice exhibits global apoptosis in embryo at ED 12.5

BLTP1^-/-^ mice are embryonic lethal. We analyzed the *BLTP1*^-/-^ embryos at multiple embryonic days (E), including E8.5 (Fig. S5A), E10.5 (Fig. S5B, C, D) and E12.5 (Fig. 5A, B, C), and found that *BLTP1*^-/-^ embryo at E12.5, but not at E8.5 or 10.5, displayed severe developmental defects (reduced size, pallor; Fig. 5B). This phenotype was similar to the embryo defect observed in VPS13A/VPS13C double KO mice, in which the defect is attributed to a defect in embryonic erythropoiesis^29^. Interestingly, the BLTP1**^-/-^** embryo did not have an evident vascular structure at embryo surface relating to the WT embryos (Fig. 5D). However, BLTP1 appears not to be required for vasculogenesis, because its depletion did not substantially impact tube formation in vitro using HUVEC cells (Fig. S5E-G). Therefore, it likely reflects a defect in embryonic erythropoiesis in BLTP1 deficient embryos.

**Fig. 5.**
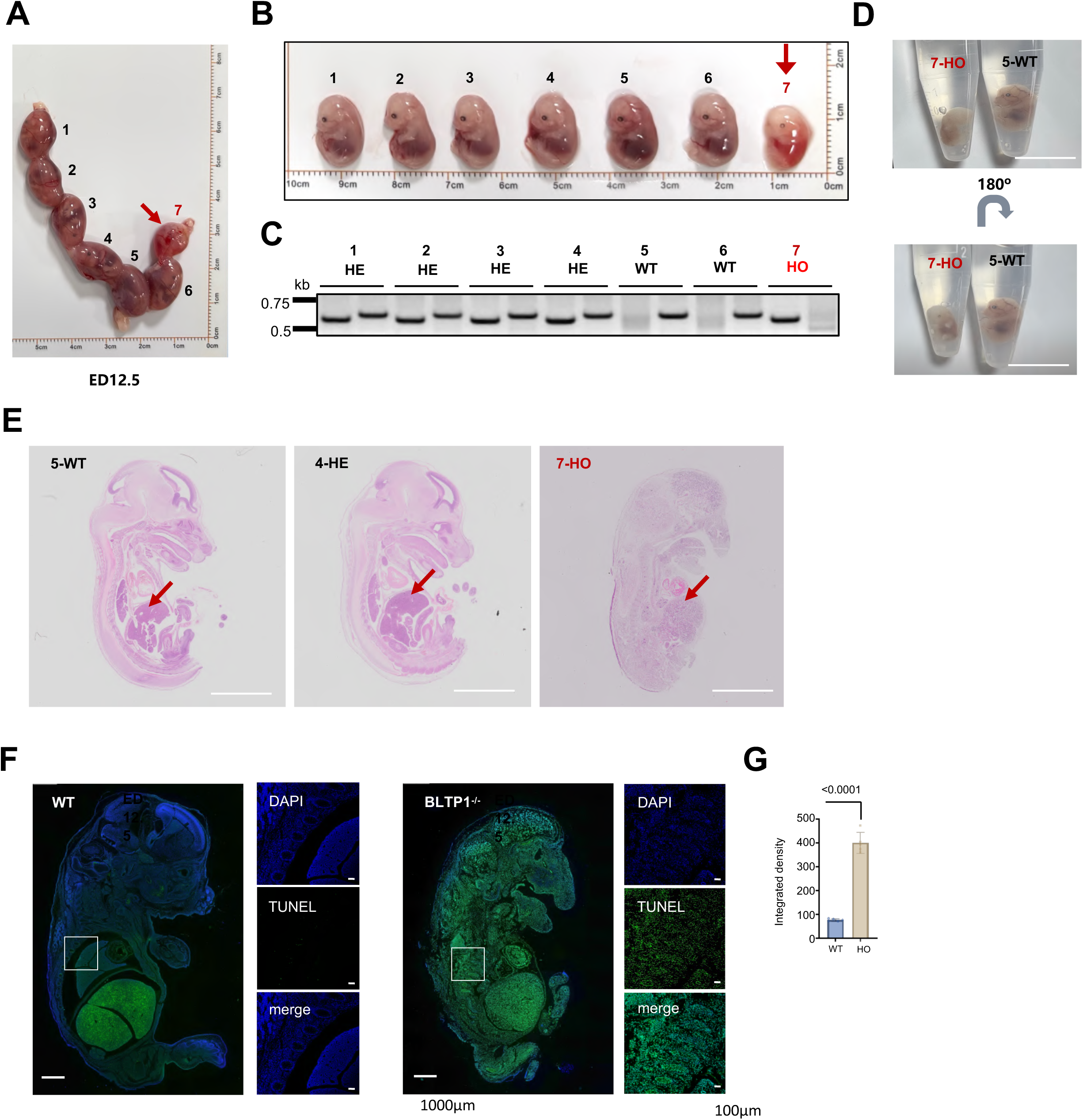
BLTP1-/- mice exhibits global apoptosis in embryo at E12.5. (A) The morphological features of uteri were observed at 12.5 days post-fertilization. A red arrow indicates an abnormal embryo. (B) Representative images of WT, heterozygous (HE), and homozygous (HO) BLTP1 KO mice embryos at E12.5. A red arrow indicates an abnormal embryo. (C) DNA gel showing genotypes of these BLTP1 KO mice embryos as shown in (B). (D) Representative images of WT (the fifth embryo; 5-WT) and HO BLTP1 KO mice embryos (the seventh embryo; 7-HO). (E) Representative images of 5-WT, 4-HE, and 7-HO embryos at E12.5 as in (B) were analyzed by hematoxylin and eosin (H&E) staining. Red arrows denote the liver. (F) Representative images of WT and BLTP1-/- embryos stained with TUNEL. (G) Quantification the TUNEL fluorescence intensity of the WT and the HO BLTP1 KO mice embryos as in (F). Three embryos from each group were quantified. Two-tailed unpaired Student’s t test. Mean ± SD. Scale bar, 500 μm in (E) and 1000 μm in the whole cell images and 100 μm in the insets in (F).

Since embryonic erythropoiesis primarily occurs in the liver, we examined E12.5 embryos by histological staining. Compared to WT littermates, the BLTP1^-/-^ embryos exhibited disorganization of multiple organs, including the brain, heart, and liver (Fig. 5E), suggesting a fundamental role for BLTP1 in embryonic development. In contrast, BLTP1^+/-^ embryos did not show substantial morphological defects (Fig. 5E). This haplosufficiency may be attributed to a compensatory mechanism involving redundancy in BLTP-mediated lipid flux and rewiring of phospholipid biosynthesis^29, 30^.

Importantly, TUNEL staining demonstrated that BLTP1^-/-^ embryo exhibited widespread apoptotic signals throughout the embryo (Fig. 5F, G), indicating that BLTP1 loss leads to elevated apoptosis during embryonic development. This finding is consistent with our cellular data and suggests that mitochondrial dysfunction-induced apoptosis likely underlies the embryonic lethality observed in BLTP1^-/-^ mice.

### Inhibition of apoptosis alleviates developmental defects in C. elegans depleted of LPD-3/BLTP1

To investigate the link between apoptosis and developmental defect caused by BLTP1 deficiency, we performed a rescuing experiment in Caenorhabditis elegans (C. elegans) depleted with LPD-3/BLTP1 by RNAi. In C. elegans, the loss of LPD-3 causes severe germline developmental defects^15^, which could be well recapitulated in a *pgl-1(ax3122)* strain with lpd-3 silencing by RNAi in this study (Fig. 6A, B, C, D, E). These defects were significantly rescued by RNAi-mediated depletion of CED-3 (apoptosis effector)—but not by LGG-1 (LC3 homolog, autophagy), ASP-3 (non-apoptotic death effector), or Fer-1 knockdown (Ferroptosis effector; Fig. 6F, G, H).

**Fig. 6.**
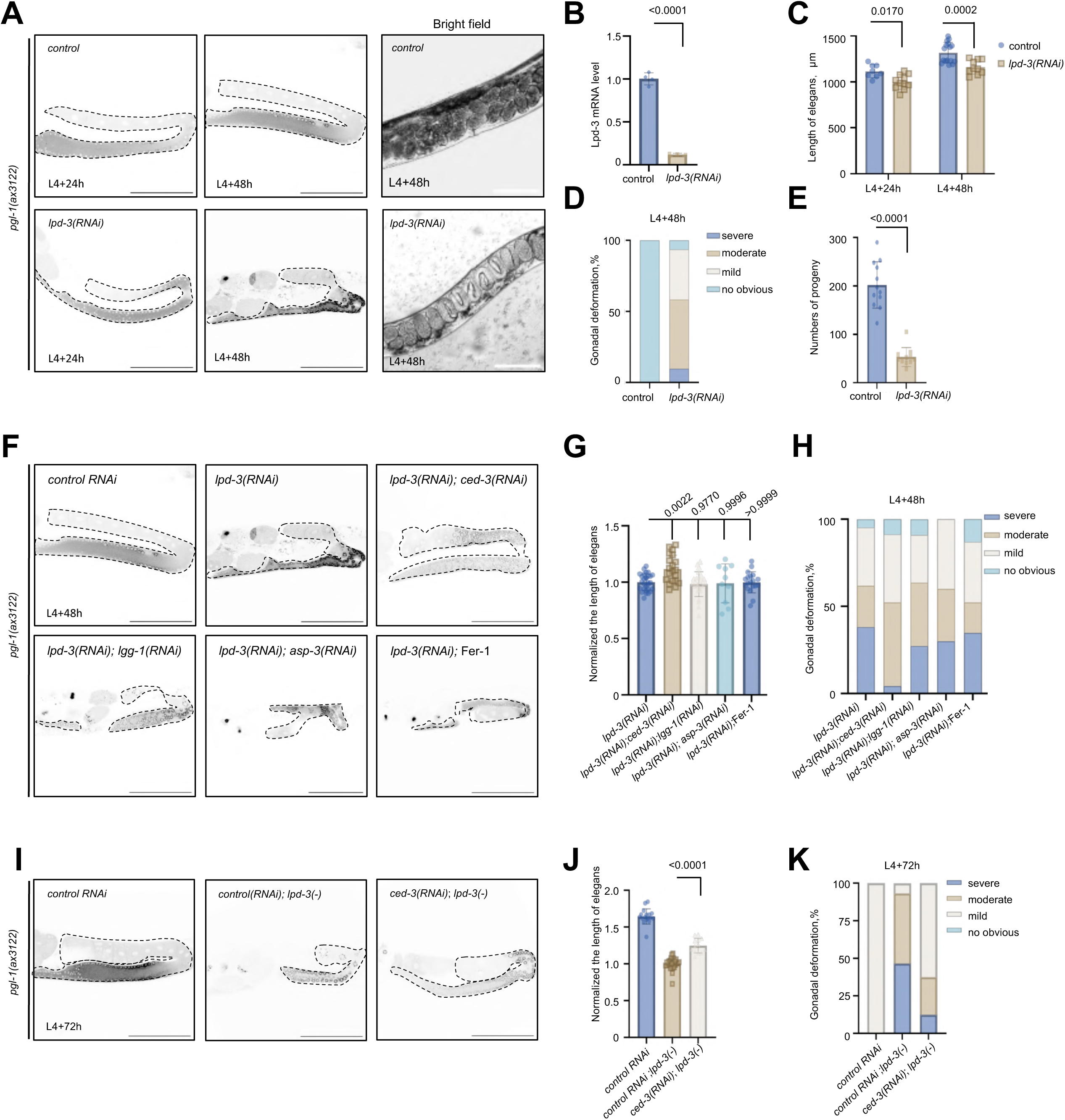
Inhibiting apoptosis mitigates gonadal deformation in LPD-3-deficient C. elegans. (A) Left: representative fluorescent images of WT or lpd-3 (RNAi) animals in the *pgl-1 (ax3122)* strain at L4+24h and L4+48h stage; right: representative bright field images of WT or lpd-3 (RNAi) in pgl-1(ax3122) strain at L4+48h stage. (B) Efficiency of lpd-3 depletion by qPCR from three independent experiments. Two-tailed unpaired Student’s t test. Mean ± SD. (C) Quantification of length of WT (n=8 and 16) and lpd-3 (RNAi) (n=10 and 10) animals at L4+24h and L4+48h stages from three independent experiments. Two-tailed unpaired Student’s t test. Mean ± SD. (D) Quantification of degree of gonadal deformation of WT (n=13) or lpd-3 (RNAi) (n=65) animals at the L4+48h stage from three independent experiments. Two-tailed unpaired Student’s t test. Mean ± SD. (E) Quantification of numbers of progeny of WT (n=12) or lpd-3 (RNAi) (n=11) animals from three independent experiments. Two-tailed unpaired Student’s t test. Mean ± SD. (F) Representative fluorescent images of control, lpd-3(RNAi), lpd-3+ced-3(RNAi), lpd-3+lgg-1(RNAi), lpd-3+asp-3(RNAi or lpd-3(RNAi)+Fer-1animals in the pgl-1(ax3122) strain. (G) Normalized length of worms as in (F). More than 10 animals from three independent experiments. Two-tailed unpaired Student’s t test. Mean ± SD. (H) Quantification of the degree of gonadal deformation in worms as in (F). More than 10 animals from three independent experiments. Two-tailed unpaired Student’s t test. Mean ± SD. (I) Representative fluorescent images of control, lpd-3(-) or lpd-3(-) +ced-3(RNAi) animals in the pgl-1(ax3122) strain. (J) Normalized length of worms as in (I). More than 10 animals from three independent experiments. Two-tailed unpaired Student’s t test. Mean ± SD. (K) Quantification of the degree of gonadal deformation of wild-type, lpd-3(-) or lpd-3(-) +ced-3(RNAi) animals. More than 10 animals from three independent experiments. Scale bar, 100 μm.

Consistently, CRISPR-Ca9-generated LPD-3 KO worms (Fig. S6A, B) exhibited gonadal deformation, particularly at late developmental stages (e.g., 72 hours post L4), and formed “wormbag” phenotypes (Fig. S6F, G). Importantly, these germline defects were substantially suppressed by CED-3 RNAi (Fig. 6I-K). These results indicate that elevated apoptosis contributes to the germline impairment caused by LPD-3/BLTP1 deficiency.

### BLTP1 depletion causes mitochondrial phospholipid accumulation

Next, we investigated whether BLTP1 deficiency drives apoptosis through mitochondrial dysfunction. As BLTP1 is a putative lipid transfer protein associated with mitochondria, we analyzed the abundance of lipid species of mitochondria by targeted lipidomics using LC-MS/MS. Following confirmation of mitochondrial fraction purity by immunoblotting (Fig. 7A), we found that BLTP1 depletion in U2OS cells markedly increased the levels of most glycerophospholipids—including PA, PG, CL, PS, PI, and PE-O—in mitochondrial fractions (Fig. 7B-I). In contrast, PE levels were largely unchanged (Fig. 7J).

**Fig. 7.**
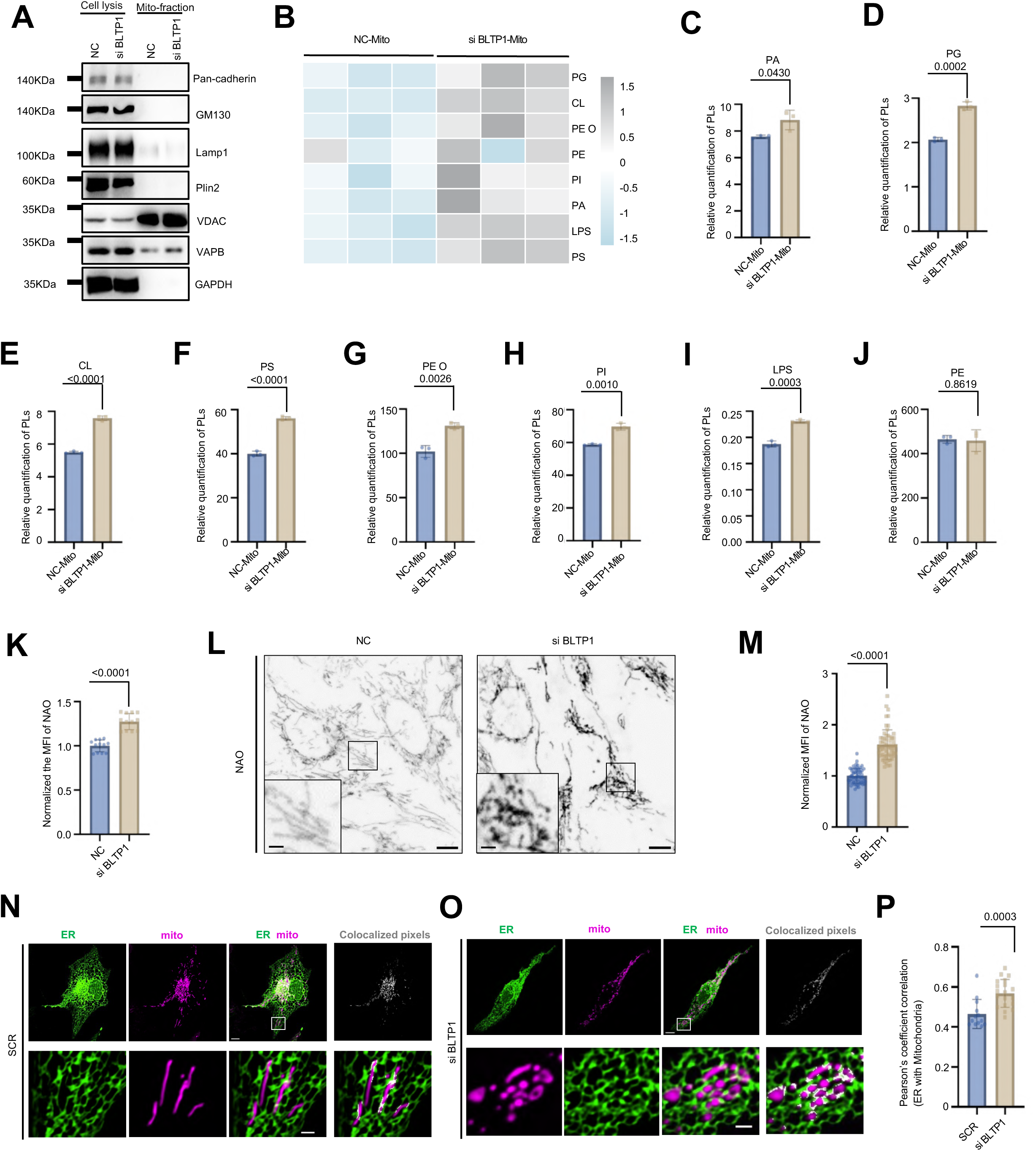
BLTP1 depletion causes mitochondrial phospholipid accumulation. (A) Representative western blots showing the specificity of the fractions enriching mitochondrial membranes from control or BLTP1-depleted U2OS cells. (B) Lipidomics of phospholipids in control or siRNA-mediated BLTP1-depleted U2OS cells. n = 3 independent experiments. (C-J) The relative quantification of phospholipids in control or BLTP1-depleted cells. n = 3 independent experiments. (K) Flow cytometry analyses of NAO fluorescence intensity of control (n=10000) or BLTP1-depleted (n=10000) U2OS cells from three independent experiments. Two-tailed unpaired Student’s t test. Mean ± SD. (L) Representative confocal images of control or BLTP1-depleted U2OS cells stained with NAO with insets. (M) Normalized NAO fluorescence intensity of control (n=60) and or BLTP1-depleted (n=60) U2OS cells as in (L) from three independent experiments. Two-tailed unpaired Student’s t test. Mean ± SD. (N, O) Representative confocal images of live U2OS cells expressing ER-GFP (green) and mitoBFP (magenta) treated with scrambled (N) or BLTP1 siRNA (O) with insets. Co-localized denote potential ER-mitochondrial MCSs. (P) Pearson’s correlation coefficient between the ER and mitochondria in cells as shown in (N, O). More than 13 cells were analyzed from three independent experiments. Two-tailed unpaired Student’s t test. Mean ± SD. Scale bar, 10 μm in the whole cell images and 2 μm in the insets in (L, N).

Consistent with the lipidomics result, the mitochondrial cardiolipin level was significantly elevated in BLTP1-depleted cells, as confirmed by both flow cytometry (using NAO fluorescence; Fig. 7K) and confocal microscopy (Fig. 7L, M).

The aberrant accumulation of phospholipids in mitochondria may result from impaired ER-mitochondria contact. To test this possibility, we assessed their physical interactions by high-resolution microscopy. Coloclaization analyses revealed that BLTP1 depletion did not reduce ER-mitochondria interaction but, rather, induced a moderate increase in their association (Fig. 7N, O, P), which may be attributed to mitochondria fragmentation in BLTP1-depleted cells. Our findings rule out impaired contact as the cause of lipid accumulation and instead point to a direct role for BLTP1 in mitochondrial phospholipid efflux, linking its loss to phospholipid overload and subsequent apoptosis.

### Suppressing mitochondrial PA import or CL synthesis prevents apoptosis caused by BLTP1 deficiency

To further validate the link between mitochondrial phospholipid overload and apoptosis, we performed candidate RNAi screen targeting mitochondrial factors involved in lipid transfer and metabolism (Fig. 8A). In this screening, we sought to identify suppressors of BLTP1 loss triggered apoptosis using the flow cytometry-based 7-AAD assays.

**Fig. 8.**
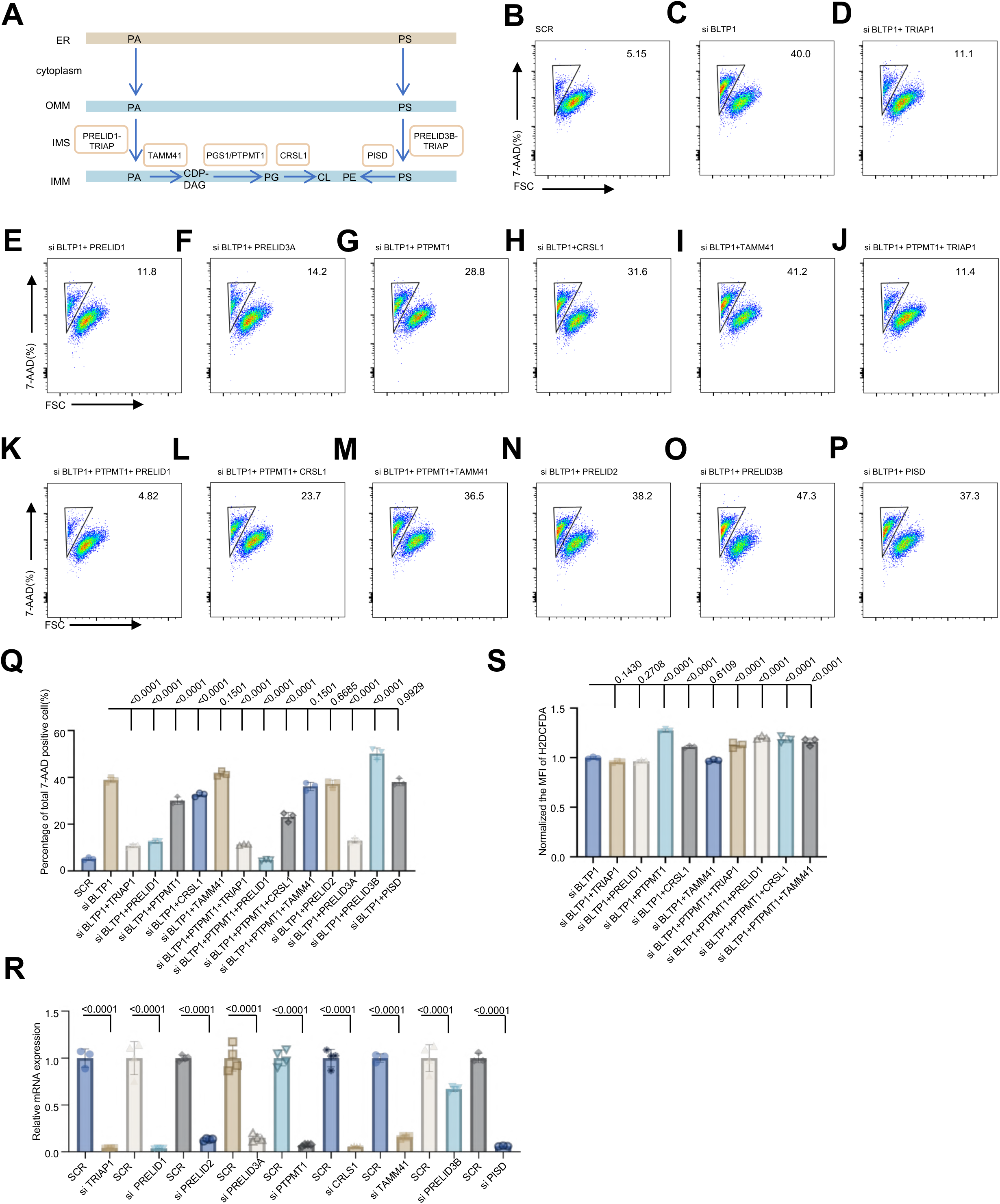
Blocking mitochondrial PA import or CL synthesis prevents apoptosis caused by BLTP1 deficiency. (A) The mammalian biosynthetic pathways of the four major classes of phospholipids (PA, PG, CL, and PE) that are biosynthesized within mitochondria. (B-P) Representative flow cytometry experiments showing percentage of 7-AAD-positive U2OS cells under control, BLTP1, lipid synthesizing enzymes or combinatorial knockdown. More than 10000 cells for each condition were analyzed. (Q) Quantification of percentage of 7-AAD-positive cells as in (B-P) in three independent assays. Two-tailed unpaired Student’s t test. Mean ± SD. (R) qPCR assays showing efficiency of siRNA-mediated suppression in cells as in (B-P). Two-tailed unpaired Student’s t test. Mean ± SD. (S) Normalized H2DCFDA fluorescence intensity of cells as in (B-P) by flow cytometry from three independent assays.

The TRIAP-PRELID complex is involved in mitochondrial PA transport to the IMM and cardiolipin synthesis^31, 32^. Intriguingly, depleting TRIAP1 (component of PA transfer complex) or its partners PRELID1/UPS1 or PRELID3A/UPS2 significantly prevented apoptosis (Fig. 8B, C, D, E, F, Q & R). Contrary to PRELID1 and PRELID3A, which have a more clearly defined function in PA transfer and cardiolipin synthesis, the role of PRELID2 and PRELID3B/UPS3 in these processes are elusive. Indeed, depletion of PRELID2 or PRELID3B/UPS3 did not have a significant rescuing effect (Fig. 8N, O, Q & R), suggesting that PRELID2 and PRELID3B/USP3 may not function in PA transport and cardiolipin synthesis.

Once synthesized in the ER, PA is shuttled from the OMM to the IMM for cardiolipin biosynthesis: PA is converted to CDP-DAG by TAMM41, which is used as a precursor for the synthesis of PG and ultimately cardiolipin, with the help of enzymes like PTPMT1, and CRLS1^33–35^. Therefore, we investigated whether suppression of these enzymes could prevent apoptotic cell death in BLTP1-depleted cells. Importantly, depletion of either PTPMT1 or CRLS1 significantly reduced the apoptotic rate compared to the control, but to a lesser extent than depletion of TRIAP-PRELID (Fig. 8G, H, Q & R).

Interestingly, suppression of TAMM41 appeared not to substantially prevent apoptosis (Fig. 8I, Q & R), possibly because CDP-DAG can be directly transferred from the ER to mitochondria rather than in situ synthesis^36^. In addition, suppression of PISD, an enzyme responsible for PE synthesis^37^, also failed to reverse the cell death caused by BLTP1 deficiency (Fig. 8P, Q & R), consistent with the lipidomic result showing no change in PE level in mitochondria in response to BLTP1 deficiency. These results thus suggest a specific link between PA import/cardiolipin synthesis and apoptotic cell death in BLTP1-depleted cells.

The combinatorial silencing of TRIAP1/PRELID1 with PTPMT1, TAMM41, or CRLS1 failed to enhance the rescue of viability compared to individual knockdowns (Fig. 8J, K, L, M, Q). This genetic epistasis indicates that these factors operate in the same pathway— specifically, in mediating the influx of PA into mitochondria for subsequent CL synthesis. Suppression of this pathway is thus sufficient to prevent apoptosis induced by BLTP1 deficiency.

Notably, despite the significant suppression of apoptosis upon blockade of PA flux or CL synthesis, the concomitant elevation in cellular ROS remained largely unchanged, as shown by flow cytometry analyses (Fig. 8S). These findings support a model where cellular ROS contributes to, but is not the sole executor of, the apoptotic cell death following BLTP1 loss. We therefore conclude that BLTP1 maintains mitochondrial lipid homeostasis by counterbalancing the influx of lipids required for CL synthesis.

### BLTP1 binds phospholipids with preference for PG and PA under mitochondrial stress

It is intriguing that BLTP1 deficiency led to mitochondrial phospholipid overload, suggessting its role in phospholipid efflux out of mitochondria. This idea prompts us to identify lipid species bound by endogenous BLTP1^Flag using using liquid chromatography-tandem mass spectrometry (LC-MS/MS) in U2OS cells, with rigorous washes of the protein before lipid analysis, according to the protocol used in our previous study^38^. BLTP1^Flag mainly associated with glycerophospholipids including PC, PE, and PS (Fig. 9A), and did not show strong lipid binding preference, similar with other BLTP proteins^39–41^.

**Fig. 9.**
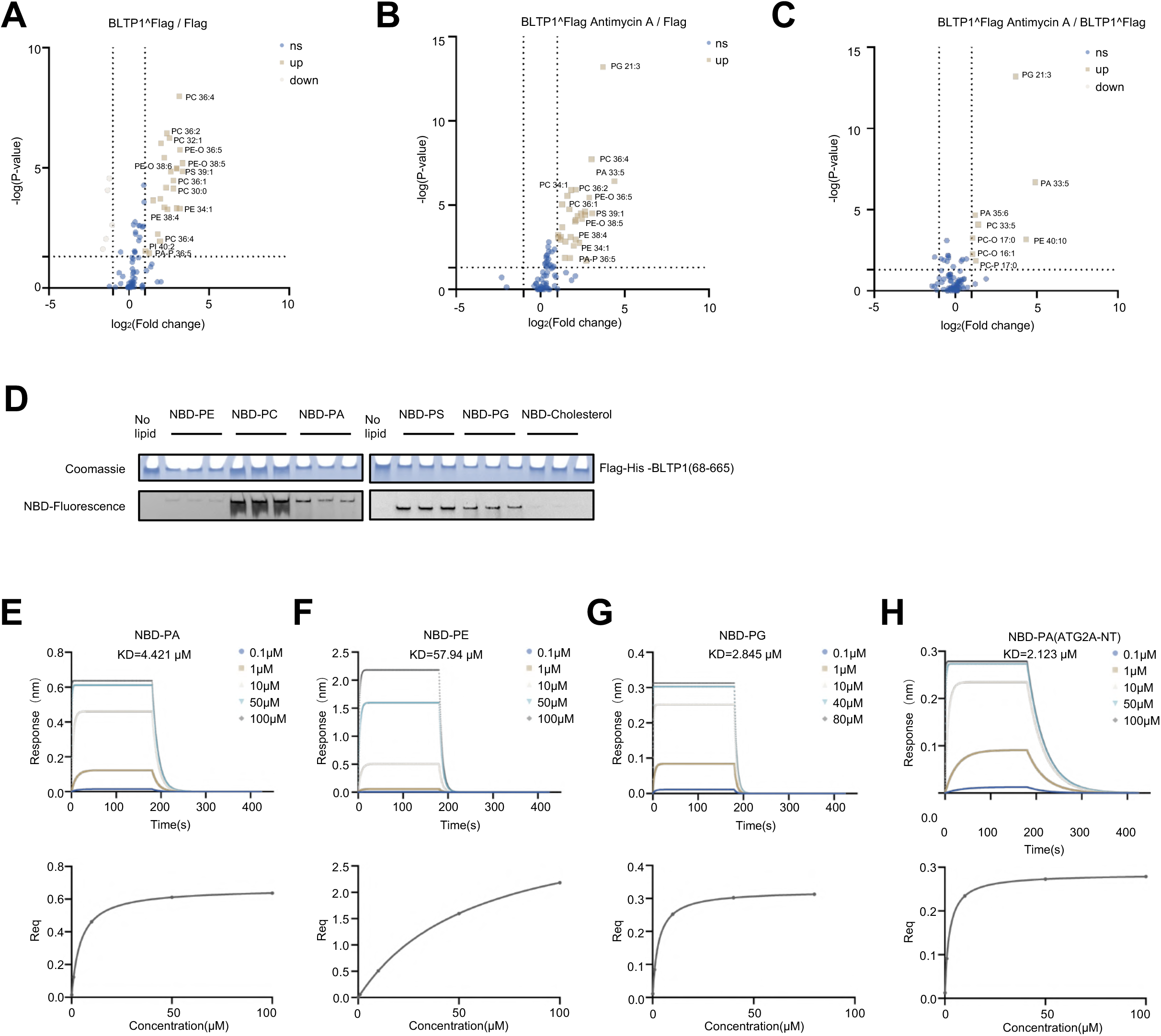
BLTP1 binds phospholipids with preference for PG under mitochondrial stress. (A) Volcano plot of phospholipids associated with BLTP1^Flag in the knock-in cell line compared with phospholipids associated with vector only. Candidates that were considered significant (−log [P value] > 1.3; P < 0.05) were labeled in light brown (Log2 [fold change] > 1; increased in abundance) or gray (Log2 [fold change] < -1 decreased in abundance). (B) As in (A), volcano plot of phospholipids associated with BLTP1^Flag in presence of Antimycin A compared with phospholipids associated with vector only. (C) As in (A), volcano plot of phospholipids associated with BLTP1^Flag in presence vs absence of Antimycin A. (D) Representative in vitro lipid-protein co-migration assays for purified BLTP1-NT (residues 68-665) fragment. (E-H) BLI assays showing the binding affinity of BLTP1-NT-(68-665) protein (0.2 μg/uL) with PA (E), PE (F), PG (G) at different concentrations. ATG2A (1-380) fragment (0.2 μg/uL) was tested as a control (H).

Remarkably, when mitochondria were stressed by Antimycin A, BLTP1^Flag could also associate with PG and PA (Fig. 9B), and the association was more evident when analyses were performed with vs without Antimycin A (Fig. 9C).

We next investigated whether BLTP1 bound these phospholipids using in vitro protein-lipids co-migrating assays. Purified BLTP1 fragment (resideus 68-665) was co-migrated with nitrobenzoxadiazole (NBD)-labeled PA, PG, PC, PS and PE, but not cholesterol, as assessed by native gel electrophoreses (Fig. 9D), consistent with the in vivo lipidomics.

We next assessed the lipid-binding affinity of BLTP1 using a biolayer interferometry (BLI) assay. A biotinylated fragment of BLTP1 was immobilized on streptavidin sensors and incubated with increasing concentrations of various lipids. Control experiments without the BLTP1 fragment were performed to subtract background signal from lipid-sensor interactions. BLI measurements yielded dissociation constants (Kd) of 2.845 μM for PG, 4.421 μM for PA, and 57.94 μM for PE (Fig. 9E–G). Among these lipids, BLTP1 exhibited the highest binding affinity toward PG, which aligns with our earlier lipidomics and protein-lipid co-migration results. Additionally, the measured Kd of PA for an ATG2A fragment (residues 1–380) was 2.123 μM (Fig. 9H), indicating a similar binding affinity between these two bridge-like lipid transport proteins (BLTPs). Interestingly, other classes of lipid transfer proteins, such as Sec14-like proteins, have been reported to exhibit dissociation constants in the nanomolar range (e.g., ∼42.54 nM)^42^, reflecting a significantly higher lipid-binding affinity than that of BLTP1 and ATG2A. This discrepancy may stem from fundamental differences in lipid transfer mechanisms: Sec14-like proteins typically operate via a shuttle mode, whereas BLTPs function through a bridge-like mode.

## Discussion

In this study, we identified the MAM protein FKBP8 as a novel BLTP1-interacting protein, and the interaction with FKBP8 was sufficient and required to recruit BLTP1 to ER-mitochondrial MCS. We demonstrate that BLTP1 deficiency resulted in phospholipid overload, accompanied by increased mitochondrial ROS, bioenergetic dysfunction, and apoptosis. Crucially, inhibiting enzymes within the CL synthesis pathway (e.g., PTPMT1, CRLS1) or mitochondrial lipid transfer proteins (e.g., PRELID1) rescues apoptosis in BLTP1-deficient cells. Collectively, these findings establish that BLTP1 mediates the essential transfer of phospholipids out of mitochondria to maintain mitochondrial function and cellular viability. Our results challenge prevailing models of BLTP function and illuminate novel mechanisms governing inter-organellar lipid flux (model; Fig. S7).

The canonical view posits that BLTP family proteins transfer lipids from the ER (the primary site of phospholipid synthesis) towards peripheral organelles, with lipid flow proceeding directionally from the N-terminus to the C-terminus of the protein. Our findings fundamentally challenge this paradigm. We demonstrate that BLTP1 operates in the reverse direction, extracting phospholipids (specifically PG, PA and likely their derivatives) from mitochondria and transferring them to the ER. This mitochondrial-to-ER lipid flow aligns with the directionality inferred from structural predictions for BLTP1 and, critically, is consistent with functional data for its yeast orthologue, Csf1, which transfers PE from mitochondria to the ER^15^. Our data thus support a model where lipid movement through BLTP1 occurs from the C-terminus (facing mitochondria) to the N-terminus (facing the ER). Despite we can not rule out the possibility that BLTP1 is able to transfer lipids bi-directionally in response to different cellular contexts, this revised model necessitates a broader re-evaluation of lipid flow directionality for other BLTP family members and underscores the context-dependent specialization of these transporters at distinct membrane contact sites.

While ER-mitochondria contact sites are established hubs for phospholipid exchange, the specific LTPs responsible for shuttling phospholipids between these organelles have remained elusive. Our identification of BLTP1, recruited via FKBP8, as the mediator of mitochondrial phospholipid export to the ER fills a significant gap in this fundamental process. The FKBP8-BLTP1 axis represents a defined molecular tether and transport machinery at these contact sites. The severe lipid dyshomeostasis and functional collapse observed upon BLTP1 loss underscore its non-redundant role in maintaining mitochondrial lipid equilibrium. This discovery provides a crucial foundation for future investigations into the composition, regulation, and dynamics of ER-mitochondria contact sites. It opens avenues to explore how BLTP1 interacts with or complements other putative LTPs at this interface and how its activity is modulated in response to metabolic or stress signals.

The directionality of BLTP1-mediated transport prompts the question: what is the functional significance of exporting phospholipids from mitochondria? Our data and emerging literature suggest several critical roles: 1) ER-Requiring Biosynthesis: As demonstrated for yeast Csf1 transferring mitochondrial-derived PE, a primary function is likely to supply the ER with specific phospholipid precursors essential for biosynthetic pathways. PE is a direct component required for GPI-anchor biosynthesis in the ER. BLTP1-mediated export of PE (and potentially other lipids) could similarly fuel GPI-anchoring or other ER-dependent modification pathways in mammals. 2) While yeast studies suggest Csf1-derived PE is dispensable for bulk autophagy under normal conditions (relying on ER-derived PE via the Kennedy pathway), a critical role for BLTP1 may emerge during metabolic constraint. If ER-derived PE synthesis is impaired (e.g., Kennedy pathway inhibition) or exogenous PE uptake is limited, BLTP1-mediated mitochondrial PE export could become essential for conjugating LC3 to phosphatidylethanolamine (LC3-II formation) and sustaining autophagic flux. This potential “backup” role warrants investigation in stress or disease models. 3) Our observation of PG accumulation in BLTP1-deficient mitochondria suggests a third key function: supplying PG to the endolysosomal system. PG exported via BLTP1 could serve as a precursor for the in situ synthesis of Bis(monoacylglycero)phosphate (BMP), also known as Lysobisphosphatidic acid (LBPA), within late endosomes/lysosomes. BMP/LBPA is a crucial lipid for endosomal sorting, intraluminal vesicle formation, and cholesterol trafficking. Mitochondria-derived PG, transported by BLTP1, could thus be a vital source for maintaining endolysosomal function and cellular lipid distribution.

In summary, we establish BLTP1 as an essential phospholipid exporter at ER-mitochondria contact sites, maintaining mitochondrial lipid homeostasis by counterbalancing the influx of lipids required for CL synthesis. Its deficiency triggers a cascade of mitochondrial lipid overload, oxidative stress, dysfunction, and cell death. The reversal of the traditional BLTP transport direction paradigm highlights the dynamic and bidirectional nature of interorganellar lipid traffic. Future research should focus on the precise structural determinants of BLTP1’s lipid selection and transport mechanism, the regulatory networks controlling its activity at contact sites (including potential roles for FKBP8 beyond recruitment), and the pathophysiological consequences of disrupted mitochondrial lipid export in Alkuraya-Kucinskas syndrome linked to mitochondrial or lipid dyshomeostasis. Understanding how BLTP1 integrates mitochondrial lipid metabolism with broader cellular homeostasis will be crucial for developing targeted therapeutic strategies.

## Materials and methods

### Cell culture, transfection, RNAi

Human osteosarcoma U2OS cells (ATCC), human cervical cancer HeLa cells (ATCC), human embryonic kidney 293T (ATCC), were grown in DMEM (Invitrogen) supplemented with 10% fetal bovine serum (Gibco). All of the cell lines used in this study were confirmed free of mycoplasma contamination.

For plasmid transfection, cells were seeded at 4×10^5^ cells per well in a six-well dish∼16 h before transfection.Plasmid transfections were performed in OPTI-MEM (Invitrogen) with 4 µL PEI to each µg plasmid for 5 h, followed by trypsinization and replating onto glass-bottom confocal dishes at ∼3.5 × 10^5^ cells per well. Cells were imaged in live-cell medium (DMEM with 10% FBS and 20 mM Hepes without Penicillin and Streptomycin) ∼16–24 h after transfection.For all transfection experiments in this study, the following amounts of DNA were used per 3.5 cm well (individually or combined for co-transfection):1000 ng for BLTP1-GFP or its truncations ; 300 ng for mCherry-FKBP8 or its truncations; 300 ng for mito-BFP, Halo/mCh-Sec61β or mCh-ACSL3.

For siRNA transfections, cells were plated on 3.5 cm dishes at 30–40% density, and 2 µL Lipofectamine RNAimax (Invitrogen) and 50 ng siRNA were used per well. At 48 h after transfection, a second round of transfection was performed with 50 ng siRNAs. Cells were analyzed 24 h after the second transfection for suppression.

### Plasmids

BLTP1 (NM_001384125.1) was synthesized (Synbiob). FKBP8 (NM_001308373.2) was cloned from 293T cDNA. The ORFs of BLTP1 was cloned into mEGFP-N1 (addgene 54767) vector between SalI and SacII. FKBP8 were cloned into mCherry-C1 vector between HindIII and BamHI. 3xFlag-6xHis-BLTP1(residues 68-665) was cloned into 3xFlag-C1 vector between HindIII and BamHI. mCherry-C1 and 3xFlag-C1 were constructed by replacing the GFP tag in mEGFP-C1 (Addgene 54579) vector with mCherry or 3xFlag via AgeI and NotI.14xHis-NEDD8-FKBP8-ΔTM was cloned into 14xHis-NEDD8 vector between BamHI and HindIII.

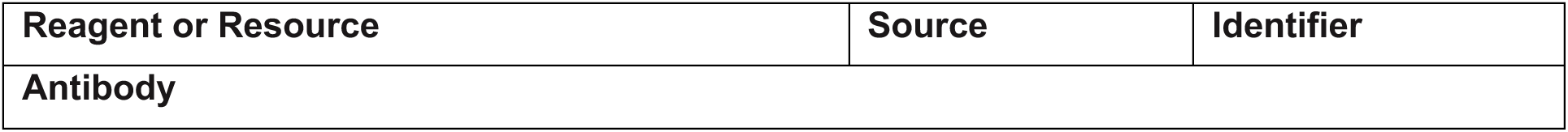
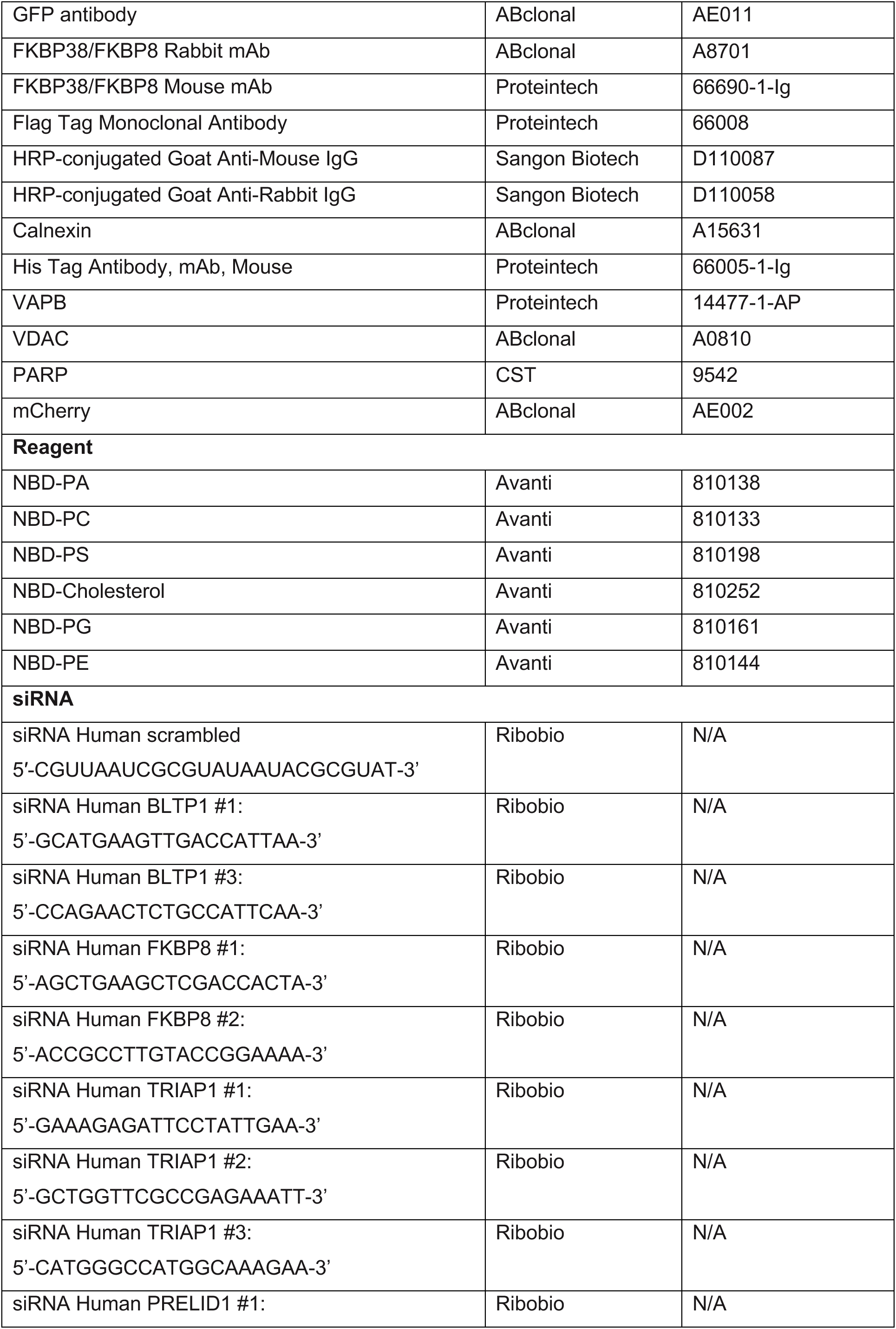
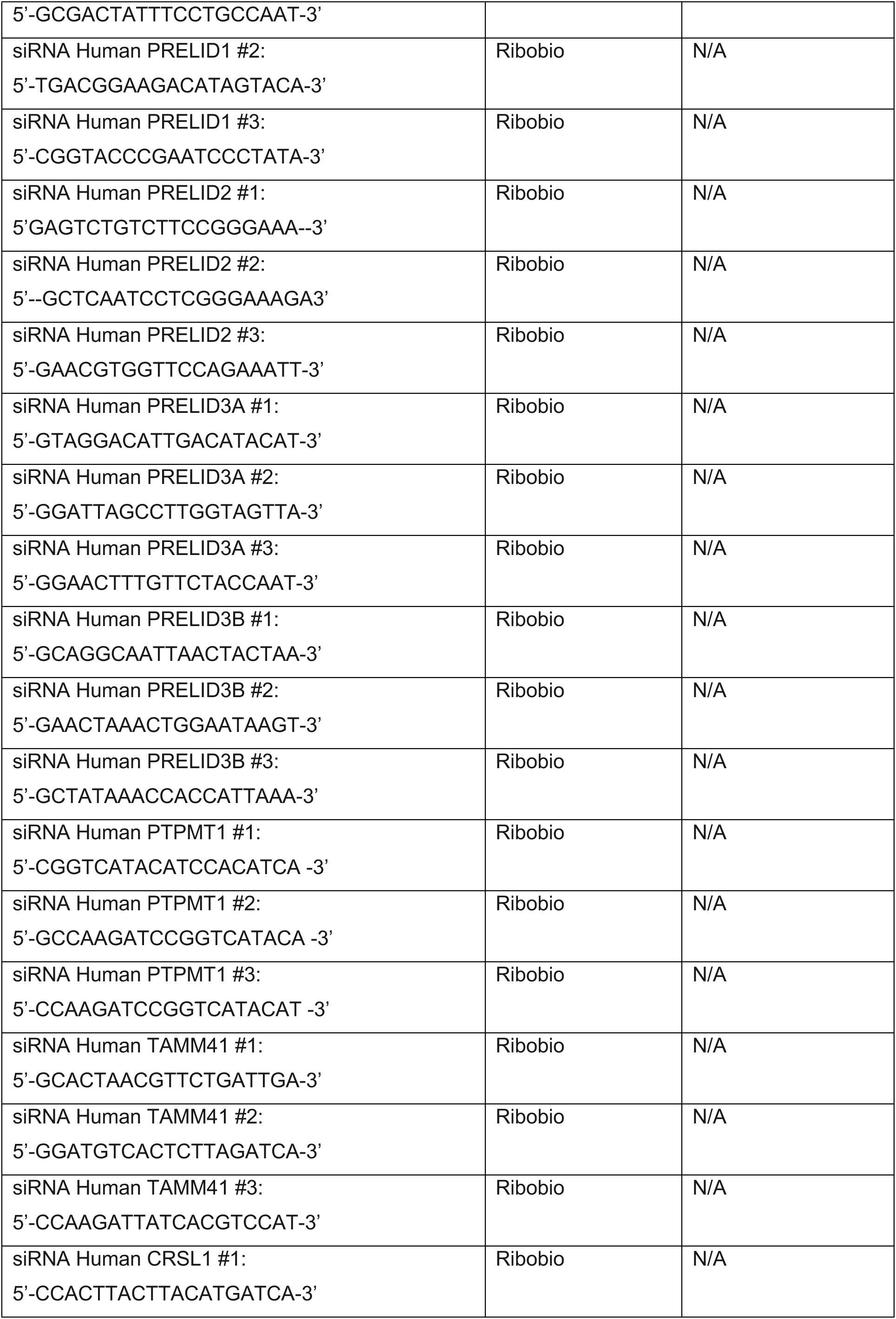
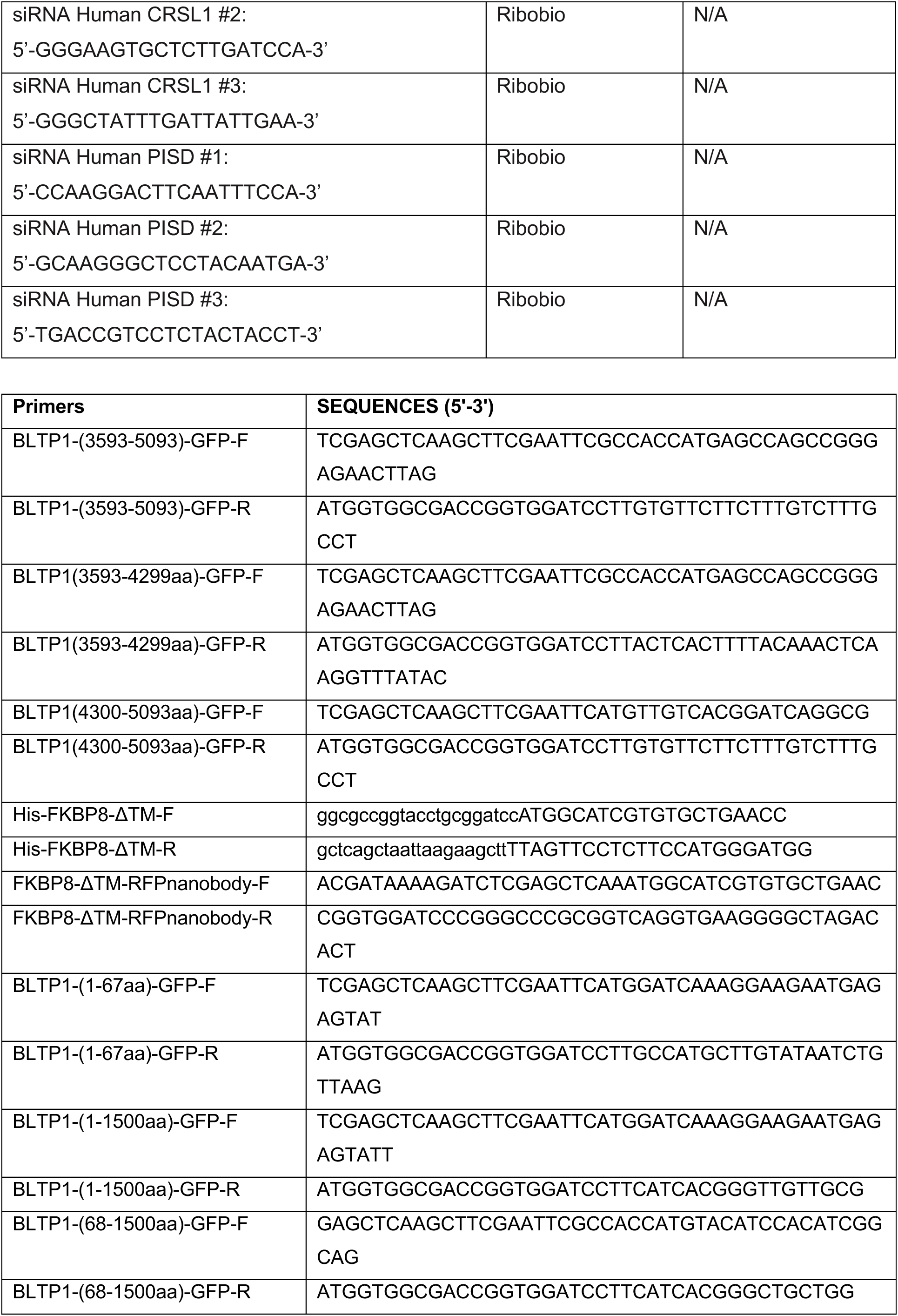
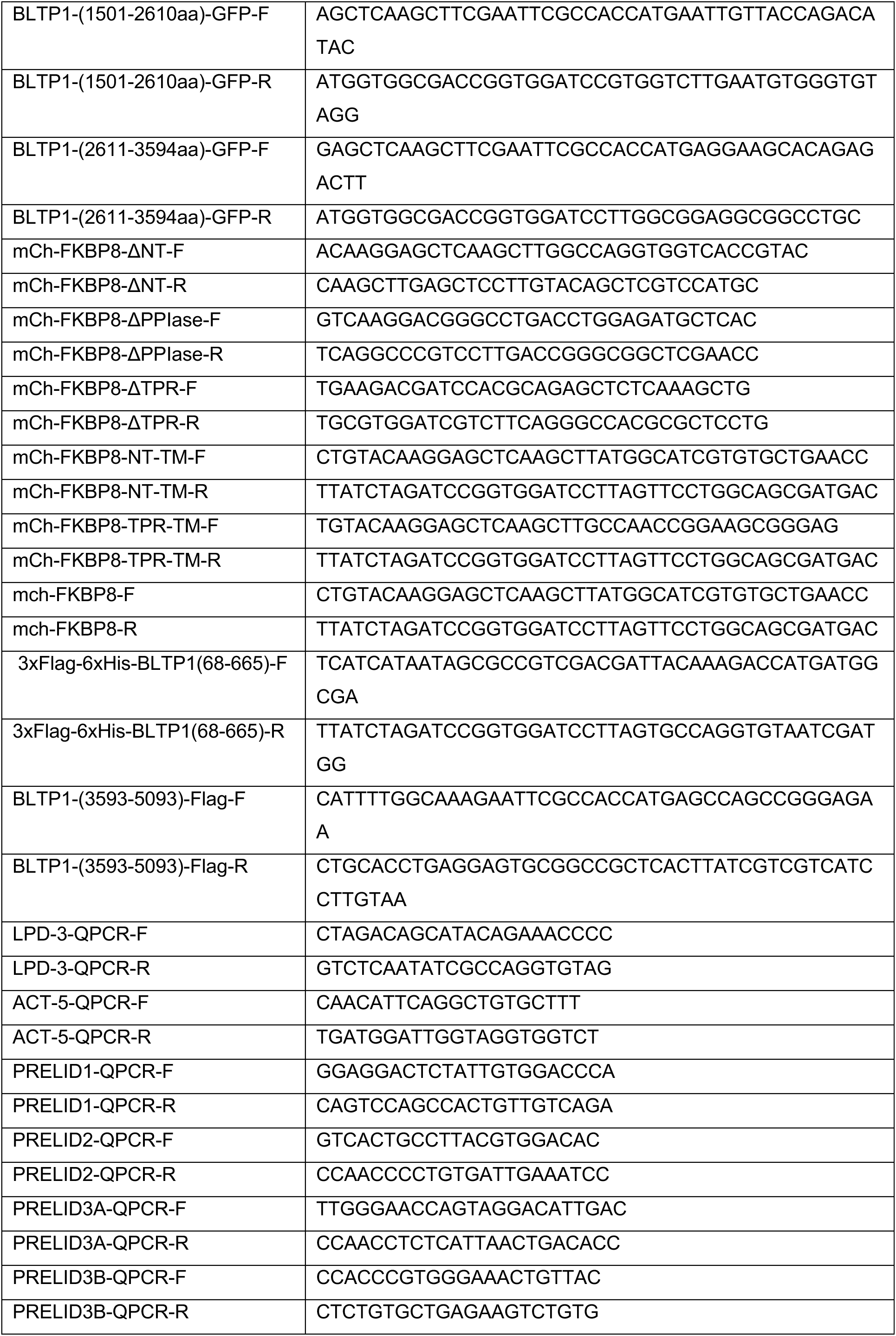
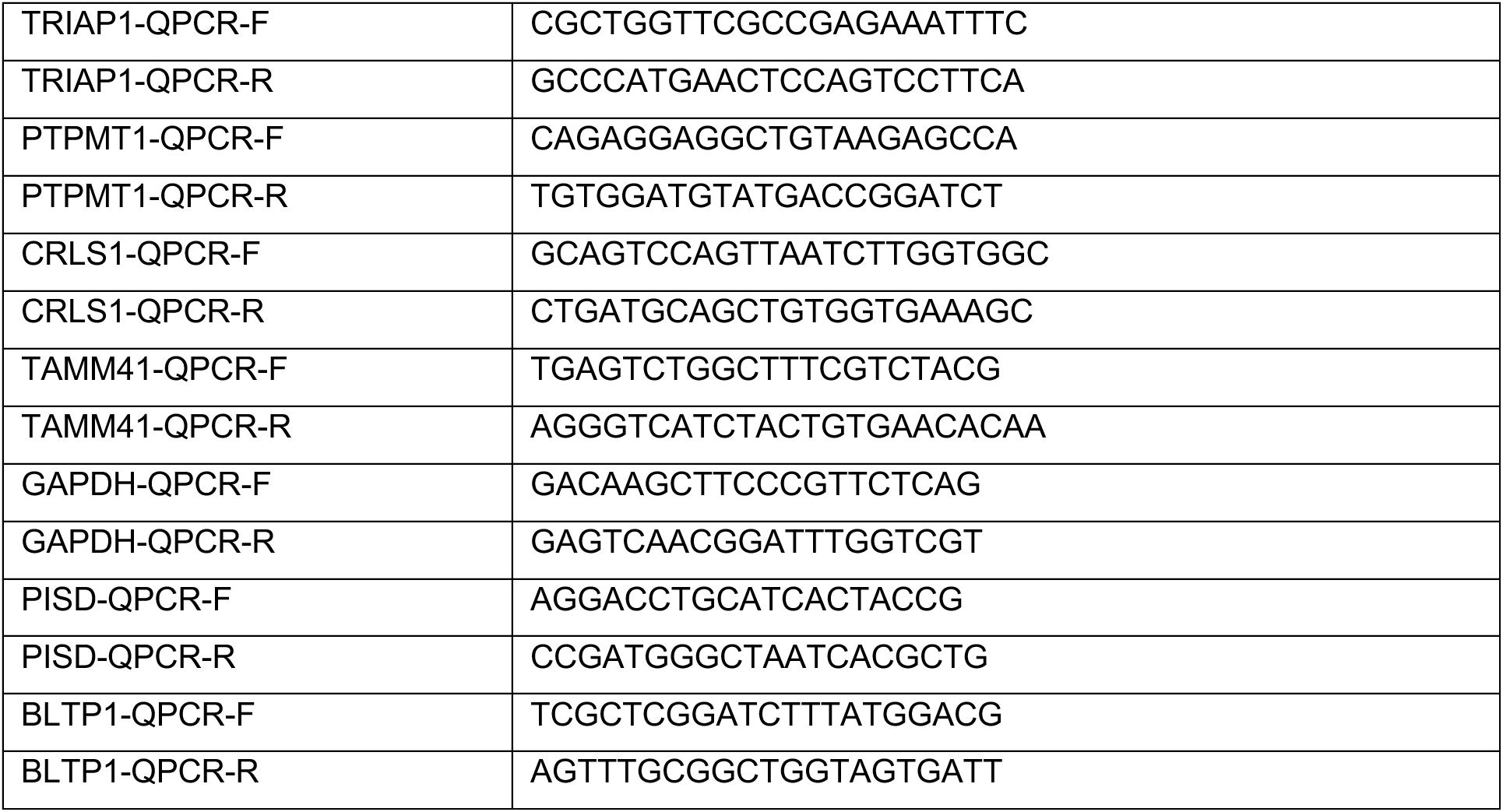

### CRISPR-Cas9-mediated gene editing

To construct BLTP1 KO cell lines, two gRNAs (5’-AGAATGGGAAAACAAATCAG-3’ and 5’-CTTCCCCAACCGATTGATTG-3’) were used.Complementary gRNAs were annealed and subcloned into the pSpCas9(BB)-2A-GFP (pX-458) vector (#48138; Addgene) between BbsI endonuclease restriction sites. Upon transfection, HeLa cells were grown in an antibiotic-free medium for 48 h, followed by single-cell sorting by fluorescence-based flow cytometry.

To construct BLTP1 KI cell lines, the CRISPR-Cas9 system was employed. Cas9 protein, sgRNA (5’-TAAGACTTCCTCTAGAACTG-3’ and 5’-TCTAGAGGAAGTCTTAATGG-3’), and the homologous repair template Oligo were transfected into U2OS cells via electroporation. Following monoclonal expansion and genotyping, the BLTP1-3×Flag cell line was successfully established. Positive monoclonal cell lines were confirmed by Sanger sequencing of PCR amplicons.

### Quantitative RT-PCR

Cells were transfected with either scrambled or BLTP1 siRNA/shRNA. 4 days after transfection, RNA was isolated with Trizol (ThermoFisher), and cDNA was reversely transcribed using RverTra Ace (TRT-101, TOYOBO) according to the manufacturer’s instructions. The cDNA was analyzed using quantitative PCR with qPCR Mix (QPS-201, TOYOBO) using the following primers for quantifications of mRNA levels.

### Western Blotting

After being heated to 95°C for 10 min, the SDS sampling buffer was used to lyse the cells. The resulting samples were subjected to SDS-PAGE electrophoresis and transferred to a Nitrocellulose transfer membrane (HATF00010; Millipore). The membrane was then blocked with 5% (m/v) non-fat powdered milk at room temperature for 1 h and then was incubated with primary antibodies at 4°C overnight. After washing three times, membranes were incubated with secondary antibodies that were conjugated with horseradish peroxidase (1/10,000) for 1 h at room temperature. Visualization was performed using enhanced chemiluminescence (P0018M-2; Beyotime).

### GFP-trap assay

GFP trap was used for detection of protein–protein interactions and were performed according to the manufacturer’s protocol. 5% input was used in GFP traps unless otherwise indicated. Briefly, cells were lysed in Lysis buffer (50 mM Tris-Cl pH 7.5, 150 mM NaCl, 0.5 mM EDTA, 0.5 % NonidetTM P40 Substitute). Lysates were centrifuged at 20,000 g for 10 min at 4°C and remove the pellets. Supernatants were incubated with GFP-Trap agarose beads for 4 h at 4°C with gentle rocking. The beads were pelleted and washed five times with wash buffer (50 mM Tris-HCl pH 7.5, 150 mM NaCl, 0.5 mM EDTA, 1x Proteases Inhibitor cocktail), then were boiled with SDS sample buffer. Proteins of interest were analyzed by immunoblotting.

### Protein expression and purification

3xFlag-6xHis-BLTP1(residues 68-665) was expressed in Expi293 cells (Invitrogen) according to the manufacturer’s instructions for 72 h. Cells were pelleted, resuspended in a buffer containing 25 mM HEPES pH 7.4, 150 mM NaCl, 2 mM MgCl2 and protease inhibitor cocktail, and were lysed via sonication. After centrifugation at 20,000 g for 1 h, the collected supernatant was incubated with anti-Flag M2 affinity resin (Sigma) for 2 h. The resin was washed with washing buffer (25 mM HEPES pH 7.4, 150 mM NaCl, 2 mM MgCl2) and was eluted with the washing buffer supplemented with 200 μg/ml 3×FLAG peptide.

14xHis-NEDD8-FKBP8-ΔTM was transformed into Escherichia coli BL21 (DE3) cells, and were incubated at 37°C until the optical density (OD) at 600 nm reached 0.6–0.8. Then cells were induced with 1 mM IPTG at 37°C for 4 hours. His fusion proteins were purified via Ni-NTA resin (QIAGEN).

### In vitro Pull-down assays of GFP Tag and His Tag

HEK293 cells transiently transfected with BLTP1-CT-GFP were lysed in high-salt lysis buffer (RIPA buffer containing 500 mM NaCl, proteasome inhibitors and PMSF). GFP-Trap beads were used to pellet BLTP1-CT-GFP from cell lysates, followed by washing with high-salt lysis buffer for 10 times. The GFP beads were incubated with purified 14xHis-NEDD8-FKBP8-ΔTM overnight at 4°C, respectively, followed by washing beads with freshly prepared HNM buffer (20 mM HEPES, pH 7.4, 0.1 M NaCl, 5 mM MgCl2, 1 mM DTT and 0.2% NP-40). GFP beads were resuspended in 100 μL 2 x SDS-sampling buffer. Re-suspended beads were boiled for 10 min at 95°C to dissociate protein complexes from beads. Western blotting was performed using anti-GFP or His antibodies. The coomassie staining was performed for purified 14xHis-NEDD8-FKBP8-ΔTM.

### Live imaging by confocal microscopy

Cells were grown on 35mm-glass bottom confocal dishes. Confocal dishes were loaded to a laser scanning confocal microscope (LSM780, Zeiss, Germany) equipped with multiple excitation lasers (405nm, 458nm, 488nm, 514nm, 561nm, 633nm) and spectral fluorescence GaAsP array detector. Cells were imaged with the 63×1.4-NA iPlan-Apochromat 63x oil objective using the 405-nm laser for BFP, 488-nm for GFP, 561-nm for OFP, tagRFP or mCherry,and 633nm for Janilia Fluo® 646 HaloTag® Ligand. Cells were imaged in live cell chamber supplied with 5% CO2 at 37°C.

### Immunofluorescence staining

Cells were fixed with 4% PFA (paraformaldehyde, Sigma) in PBS for 10 min at room temperature. After washing with PBS three times, cells were permeabilized with 0.1% Triton X-100 in PBS for 15 min on ice. Cells were then washed three times with PBS, blocked with 3% BSA in PBST for 1 h, incubated with primary antibodies in diluted blocking buffer overnight, and washed with PBST three times. Secondary antibodies were applied for 1 h at room temperature. After washing with PBST three times, samples were mounted on Vectashield (H-1000; Vector Laboratories).

### Mass spectrometry for identification of BLTP1-interacting Proteins

The identification of endogenous BLTP1-interacting proteins by mass spectrometry (MS) was described in our previous study^38^. Briefly, the bound proteins from the BLTP1^Flag line were extracted from Flag M2 agarose beads using SDT lysis buffer (4% SDS, 100 mM DTT, 100 mM Tris-HCl, pH 8.0), followed by sample boiling for 3 min and further ultrasonicated. Undissolved beads were removed by centrifugation at 16,000 g for 15 min. The supernatant was collected. Protein digestion was performed with the FASP method. Briefly, the detergent, DTT, and IAA in the UA buffer were added to block-reduced cysteine. Finally, the protein suspension was digested with 2 µg trypsin (Promega) overnight at 37°C. The peptide was collected by centrifugation at 16,000 g for 15 min. The peptide was desalted with C18 StageTip for further LC-MS analysis. LC-MS/MS experiments were performed on a Q Exactive Plus mass spectrometer that was coupled to an Easy nLC (Thermo Fisher Scientific). The peptide was first loaded to a trap column (100 µm × 20 mm,5 µm, C18, Dr. Maisch GmbH) in buffer A (0.1% formic acid in water). Reverse-phase high-performance liquid chromatography (RP-HPLC) separation was performed using a self-packed column (75 µm × 150 mm; 3 µm ReproSil-Pur C18 beads, 120 °A; Dr. Maisch GmbH, Ammerbuch) at a flow rate of 300 nl/min. The RP-HPLC mobile phase A was 0.1% formic acid in water and B was 0.1% formic acid in 95% acetonitrile. The gradient was set as follows: 2–4% buffer B from 0 to 2 min, 4–30% buffer B from 2 to 47 min, 30–45% buffer B from 47 to 52 min, 45–90% buffer B from 52 min and to 54 min, and 90% buffer B kept until to 60 min. MS data were acquired using a data-dependent top20 method dynamically choosing the most abundant precursor ions from the survey scan (350–1,800 m/z) for HCD fragmentation. A lock mass of 445.120025 Da was used as the internal standard for mass calibration. The full MS scans were acquired at a resolution of 70,000 at m/z 200, and 17,500 at m/z 200. The maximum injection time was set to 50 ms for MS and 50 ms for MS/MS. Normalized collision energy was 27 and the isolation window was set to 1.6 Th. Dynamic exclusion duration was 60 s. The MS data were analyzed using Max Quant software version 1.6.1.0. MS data were searched against the UniProtKB human database (36,080 total entries, downloaded 2019.06.25). Trypsin was selected as the digestion enzyme. A maximum of two missed cleavage sites and a mass tolerance of 4.5 ppm for precursor ions and 20 ppm for fragment ions were defined for database search. Carbamidomethylation of cysteines was defined as a fixed modification, while acetylation of protein N-terminal and the oxidation of methionine were set as variable modifications for database searching. The database search results were filtered and exported with a <1% false discovery rate (FDR) at peptide-spectrum-matched level and protein level, respectively.

### In vitro lipid-binding assay

1 μL of either NBD-labeled PE, PS, PA, PC, PG or cholesterol (1mg/mL in methanol) was incubated with19 μL purified 3xFlag-6xHis-BLTP1 (68-665) for 2 h at 4°C. Samples were visualized on 6% native PAGE gels. NBD fluorescence was visualized using ChampChemi610 plus, and comigrated with protein was visualized with Coomassie staining.

### Mitochondria-associated ER Membranes (MAMs) Isolation

Cells were harvested and resuspended in Solution A (0.32 M Sucrose, 1 mM NaHCO₃, 1 mM MgCl₂, 0.5 mM CaCl₂, supplemented with Protease Inhibitors cocktail). The cell suspension was homogenized on ice using a Teflon pestle homogenizer operating at 4,000 RPM. Homogenization was paused every 25 strokes to monitor cell integrity by light microscopy, and the process was concluded when 80–90% of the cells were disrupted. The homogenate was diluted to a 10% (w/v) concentration with Solution A and centrifuged at 1,400 g for 10 min at 4 °C. The supernatant was collected, and the pellet was resuspended in an equal volume of Solution A, followed by an additional 5 strokes of homogenization and centrifugation at 710 × g for 10 min at 4 °C to pellet nuclei and cellular debris. The resulting supernatant was combined with the first supernatant and centrifuged at 13,800 g for 10 min at 4 °C. The pellet, enriched in mitochondria, was resuspended in Solution A, homogenized with 5 strokes, and centrifuged again under the same conditions. This wash step was repeated once. The final mitochondrial pellet was resuspended in Solution B (0.32 M Sucrose, 1 mM NaHCO₃ with Protease Inhibitors) at a concentration of 4.8 g/mL and homogenized with 6 strokes. This resuspension was then layered at the bottom of an ultracentrifuge tube and successively overlaid with 3 mL of 0.85 M sucrose, 3 mL of 1 M sucrose, and 3 mL of 1.2 M sucrose (all prepared in 1 mM NaHCO₃). The gradient was centrifuged at 82,500 g for 2 h at 4 °C. Following centrifugation, distinct bands were isolated: the 0.32/0.85 M interface (containing myelin and membrane contaminants), the 0.85/1 M interface (enriched in ER, Golgi, and plasma membranes), the 1/1.2 M interface (synaptosomes), and the pellet (crude mitochondria). The crude mitochondrial pellet was resuspended in Isolation Medium (250 mM Mannitol, 5 mM HEPES pH 7.4, 0.5 mM EGTA, 0.1% BSA, plus protease inhibitors) and further purified by centrifugation through a 30% Percoll gradient prepared in Gradient Buffer (225 mM Mannitol, 25 mM HEPES pH 7.5, 1 mM EGTA, 0.1% BSA) at 95,000 g for 30 min at 4 °C. The heavy mitochondrial fraction (lower band) and the light fraction (upper band) were carefully collected into separate tubes. The heavy fraction was diluted 1:10 with Isolation Medium and pelleted by centrifugation at 6,300 g for 10 min at 4 °C; this wash step was repeated once to obtain the purified mitochondrial fraction. The light fraction was similarly diluted and subjected to ultracentrifugation at 100,000 g for 1 h at 4 °C to pellet the mitochondrial-associated membranes (MAMs).

### Superdex 6 Increase Size-exclusion chromatography

Size-exclusion chromatography to assess protein complex formation 200 μg or 400 μg of each protein were diluted with storage buffer to a final volume of 500 μl in a 1.5 mL tube. After incubation at 4 °C for 4 h, samples were centrifuged at 20,000 rcf for 10 minutes at 4 °C and loaded onto a Superdex 6 Increase 10/300 Gl column. Size-exclusion chromatography was performed at 4 °C on an AKTA pure system at a flow rate of 0.5 ml/min. Protein elution was monitored at 280 nm, and fractions of 0.4 mL were collected. 30 μL were removed from each fraction and added to a tube containing 10 μL 4× SDS-PAGE sample buffer. Samples were heated for 10 min at 95 °C, and 30 μL were separated by 8% SDS-PAGE. Proteins were visualized by western blots.

### ROS measurement

Cells were incubated with HBSS containing 5 μM or 10 μM H2DCFDA (HY-D0940, MCE) at 37°C for 30 min in dark.Then cells were trypsonized and then washed twice with PBS, and resuspended in fresh PBS for flow cytometry analysis. For mitochondrial superoxide detection using flow cytometry, cells were incubating with 5 mM MitoSOX-red (M36008, Thermo Fisher) at 37°C for 30 min in dark. Samples were detected with excitation at 488 nm and emission at FITC channel for H2DCFDA and FL2 channel for MitoSOX-red. Flow cytometry data were processed in FlowJo. Cells were gated for live cells in SSC-A vs FSC-A plot and for singlet cells in FSC-H vs FSC-A plot. The FITC-A and PE-A histogram of singlet cells were plotted and the mean intensity of each peak was quantified. For all flow cytometry assays, at least 3 biological replications were performed.

### Flow cytometry-based cell death assay

Cells were transfected with siRNA as above. Medium was replaced 6 h after transfection. Three days after transfection, cell death was analysed using 7-AAD (BD Pharmingen, 559925) or SYTOX™ Green Nucleic Acid Stain (ThermoFisher, S7020). Flow cytometry was performed using an Novocyte 2060R flow cytometer and at least 10,000 events were analysed per sample. Data were analysed using FlowJo. Gating was applied using FSC-A and FSC-H to select singlets (the single cell population). Subsequently, FSC-A and fluorescence parameters were used to gate and sort the positive cell population. FSC-A and SSC-A was used to remove cellular debris. Notably, Fer-1 treatment caused non-specific signals in 7-AAD staining, and thus we used SYTOX™ Green instead in Fig 4.

### BLI assay

Measurement of PG, PA and PE binding with BLTP1 or ATG2A fragment protein was performed at 30 °C on a BLI instrument (Octet^®^R2). An SA sensor was chosen to immobilize purified biotin-tagged BLTP1 or ATG2A fragment protein. Different concentrations of PG, PA and PE were dried under N2 and then dissolved in PBS as analytes. Data were fitted by Octet Data Analysis 12.2 software to export binding constants.

### Tube network formation assay

24-well plates were coated with 30 µL Matrigel per well and polymerized at 37°C. Control and BLTP1 depleted HUVEC cells were seeded at a density of 5.0 × 10⁴ cells per well in medium contain 20uM Z-VAD-FMK. The plates were incubated at 37°C and 5% CO₂ for 8 hours to allow tube network formation. Subsequently, images from 3-4 random fields per well were captured using an inverted light microscope. The formed tubular structures were quantified with ImageJ software and its Angiogenesis Analyzer tool. Key parameters analyzed included total tube length, number of nodes.

### Statistical analysis

All statistical analyses and p-value determinations were performed in GraphPad Prism (8.0.1) All the error bars represent Mean ± SD. To determine p-values, ordinary one-way ANOVA with Tukey’s multiple comparisons test were performed among multiple groups and a two-tailed unpaired student t-test was performed between two groups.

## Acknowledgements

We thank Zhenrong Yang, Linfang Yang, Qing Tian and Weimin Wang (School of Basic Medicine Innovation research center, Huazhong University of Science and Technology) for assistance with imaging, flow cytometry and BLI experiments. We thank Multi-Omics Mass Spectrometry Core of Shenzhen Bay Laboratory for assistance with proteomics and lipidomics experiments.

This study was supported by National Natural Science Foundation of China (32371343, 92354304, 32270779), Shenzhen Bay Scholars Program, State Key Laboratory for diagnosis and treatment of severe zoonotic infectious disease.

The authors declare no competing interests.

Author contributions: Q. Chu and W. Ji conceived the project and designed the experiments. Q. Chu, T. Cui, X. Lu, Z. Zhou, J. Luo performed the experiments. Q. Chu, T. Cui, X. Lu, W. Chang, L. Deng and W. Ji analyzed and interpreted the data. W. Ji prepared the manuscript with inputs and approval from all authors.

## Data and materials availability

All the data and relevant materials, including reagents and primers, that supports the findings of this study are available from the corresponding author upon reasonable request.

---

**Fig. S1.**
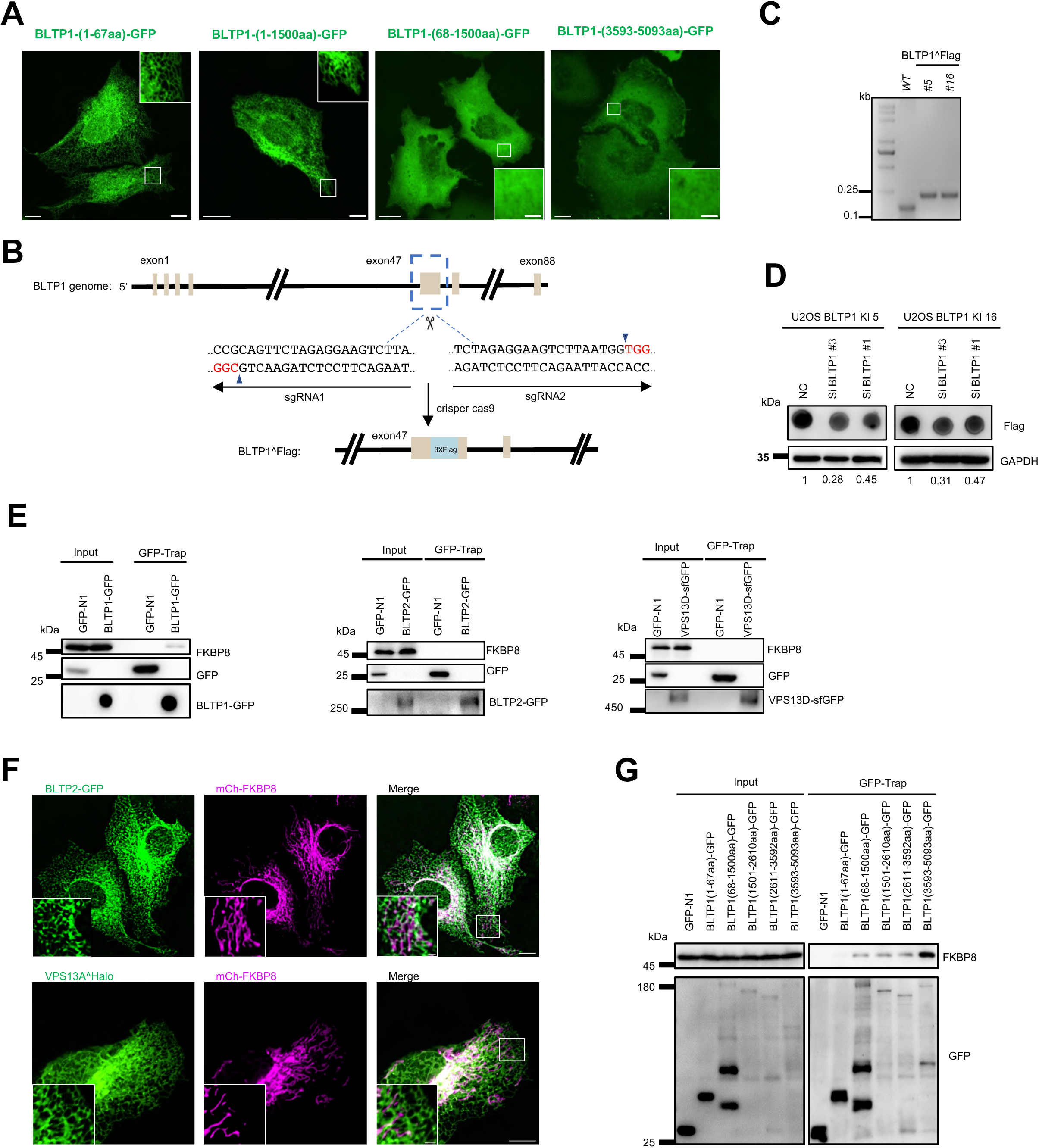
The interaction between BLTP1 and FKBP8. (A) Representative live-cell images of U2OS cells expressing BLTP1-GFP and its truncations (green) with insets. (B) Generation of the BLTP1^Flag KI U2OS cell by CRISPR-Cas9. Two sgRNAs are used with the underlined letters, indicating the PAMs (CGG for sgRNA1 and TGG for sgRNA2) for spCas9. (C) DNA gel confirming the insertion of 3x Flag tag in the genome of the BLTP1 Knock-in cell line. Two clones (#5 and #16) were verified. (D) Representative western blot confirming the level of BLTP1^Flag in control or BLTP1-depleted cells. (E) GFP-Trap assays demonstrate a specific interaction between endogenous FKBP8 and BLTP1 but not BLTP2 or VPS13D. (F) Representative live-cell images of U2OS cells expressing BLTP2-GFP (green; top) or VPS13A-Halo (green, bottom) with mch-FKBP8 (magenta) with insets. (G) Representative GFP-Trap assays demonstrating that endogenous FKBP8 preferentially interacted with BLTP1-CT. Scar bar, 10 μm in the whole cell images and 2 μm in the insets in (A, F)

**Fig. S2.**
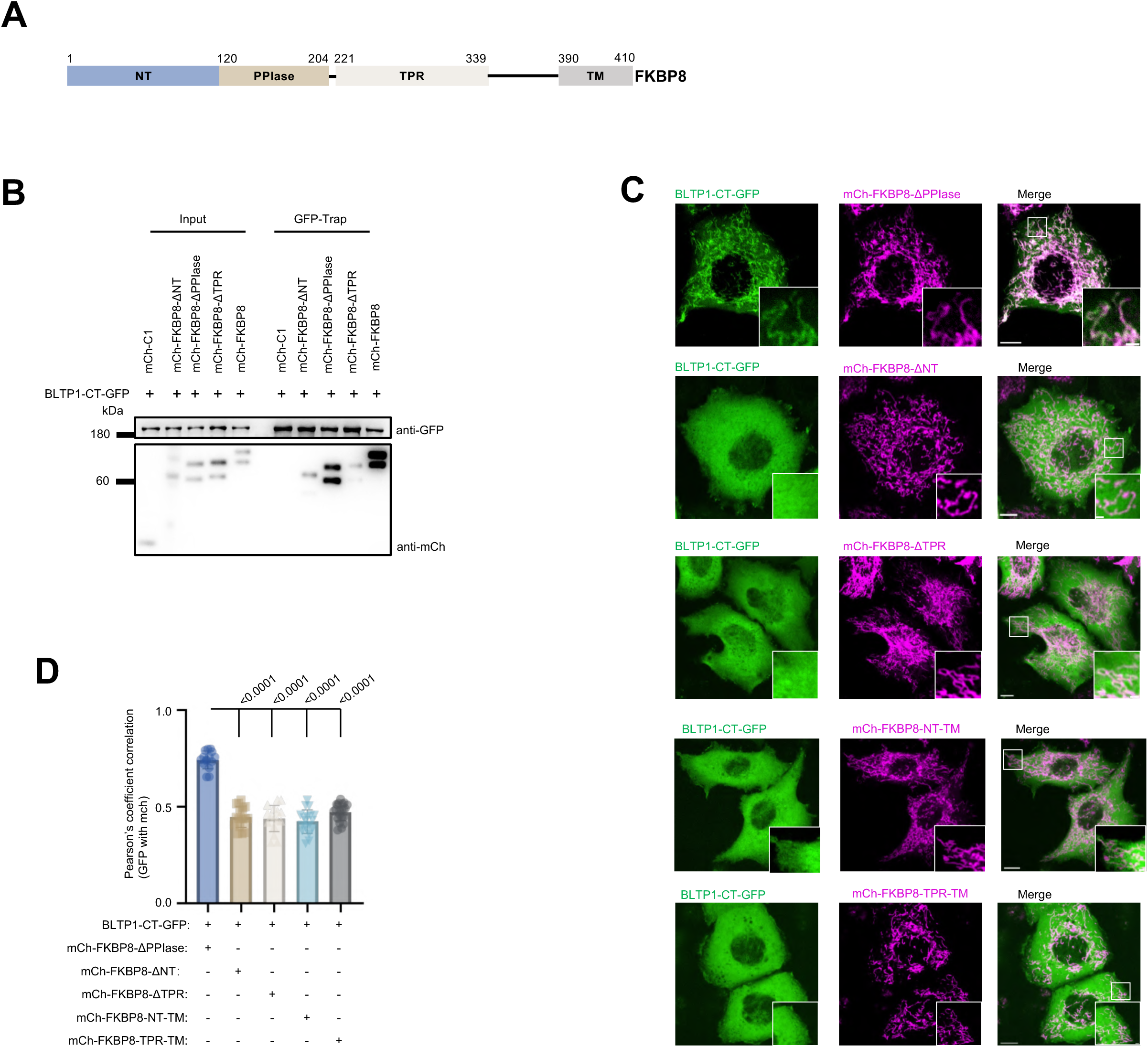
The domains of FKBP8 responsible for interacting with BLTP1. (A) The domain organization of FKBP8. (B) GFP-Trap assays showing interactions between mCh-FKBP8 truncations and BLTP1-CT-GFP in HEK293 cells. (C) Representative images of live U2OS cells expressing BLTP1-CT-GFP (green) and mch-FKBP8 truncations (magenta) with insets. (D) Pearson’s correlation coefficient in cells as shown in (C). In the quantification, more than 15 cells for each condition were analyzed from three independent experiments. Two-tailed unpaired Student’s t test. Mean ± SD. Scale bar, 10 μm in the whole cell images and 2 μm in the insets.

**Fig. S3.**
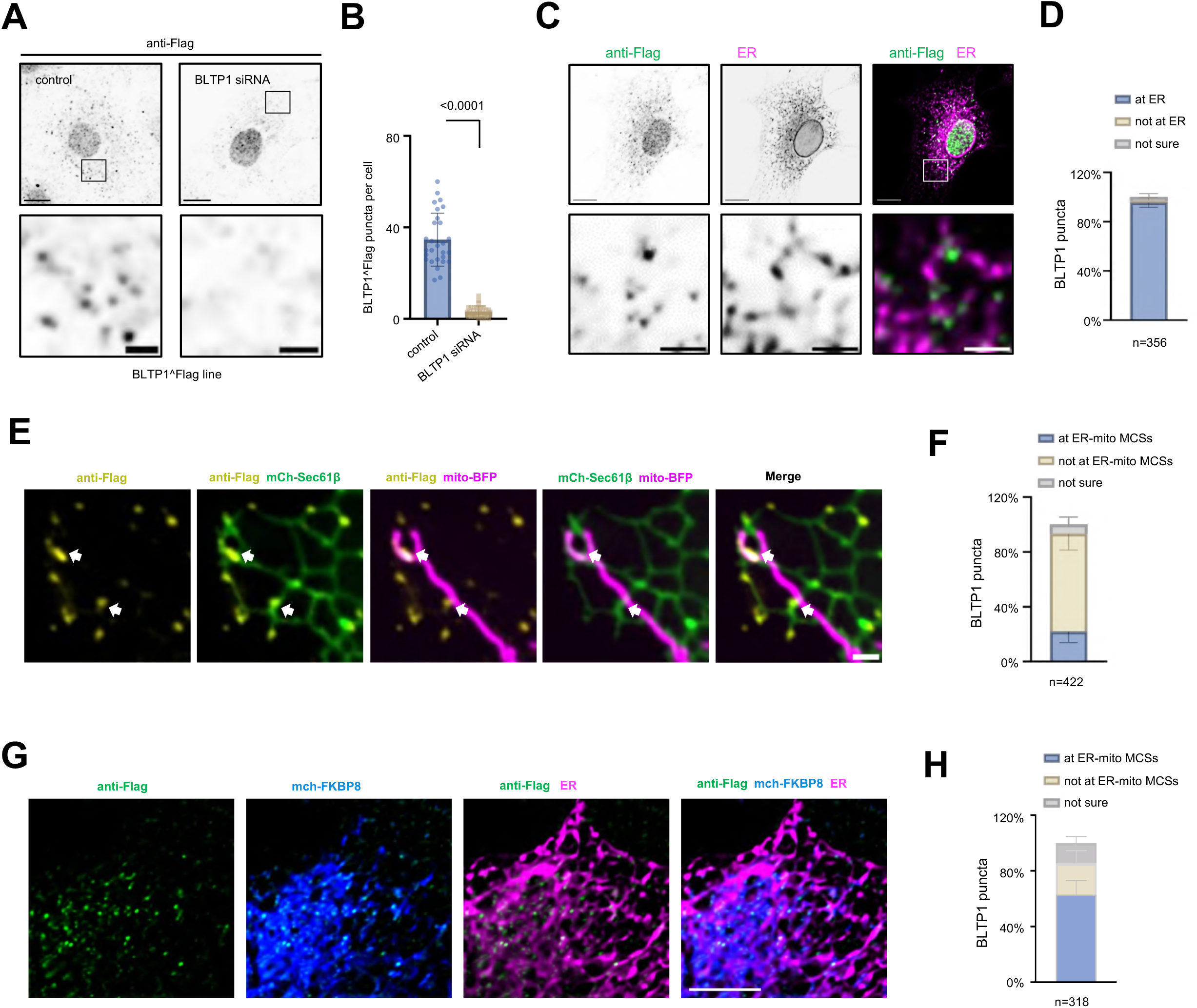
The recruitment of endogenous BLTP1 by FKBP8. (A) Confocal images of fixed BLTP1^Flag KI cells treated with scrambled (left panel) or BLTP1 siRNA (right panel) stained with antibodies against Flag. (B) Quantification of BLTP1^Flag puncta per cell as shown in (A) in three independent assays. Two-tailed unpaired Student’s t test. Mean ± SD. (C) Representative images of fixed KI cells stained with antibodies against Flag (green) and Halo-sec61β (magenta) with insets. (D) Quantification of BLTP1^Flag puncta relative to ER in cells as shown in (C). More than 10 were analyzed from three independent experiments. Two-tailed unpaired Student’s t test. Mean ± SD. (E) Representative images of fixed U2OS cells stained with antibodies against Flag (yellow) and expressing mitoBFP (magenta) and mCh-Sec61β(green) with insets. Arrows indicate BLTP1 puncta associating with mitochondria. (F) Quantification of BLTP1^Flag puncta relative to ER-mitochondrial MCSs in cells as shown in (E). More than 50 cells were analyzed from three independent experiments. Two-tailed unpaired Student’s t test. Mean ± SD. (G) Representative images of fixed U2OS cells stained with antibodies against Flag (green) and expressing mCh-FKBP8 (blue) and Halo-Sec61β (magenta) with insets. (H) Quantification of BLTP1^Flag puncta relative to ER-mitochondrial MCSs in cells as shown in (G). More than 40 were analyzed from three independent experiments. Two-tailed unpaired Student’s t test. Mean ± SD. Scale bar, 10 μm in the whole cell images and 2 μm in the insets in (A, C) and 2 μm in (E, G).

**Fig. S4.**
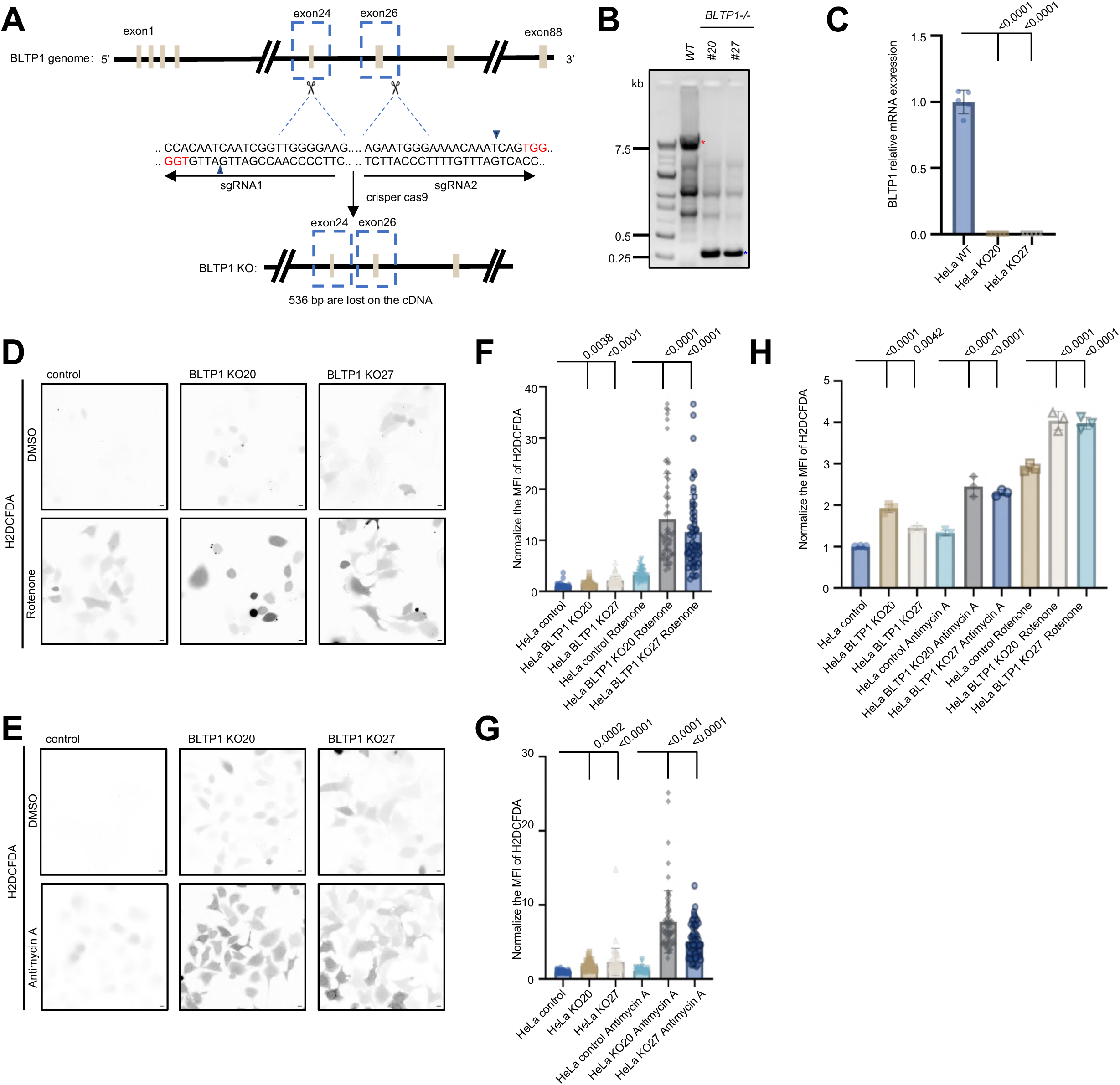
BLTP1 deficiency caused ROS elevation and apoptosis. (A) Generation of CRISPR-Cas9-mediated BLTP1 KO HeLa cell line (BLTP1-KO). Two sgRNAs are used with the underlined letters, indicating the PAMs (TGG for sgRNA1 and TGG for sgRNA2) for spCas9. (B) Genotyping of control and the BLTP1 KO cell line by DNA gel electrophoresis. (C) qPCR assays showing the efficiency of sgRNA-mediated suppression in BLTP1. (D, E) Representative images of control or two BLTP1 KO clones stained with H2DCFDA under DMSO, Rotenone (D) or Antimycin A (E) treatments. (F, G) Normalized H2DCFDA fluorescence intensity of control and BLTP1 KO cells as in (D, E). More than 60 cells were analyzed from three independent experiments. Two-tailed unpaired Student’s t test. Mean ± SD. (H) Flow cytometry analyses of H2DCFDA fluorescence intensity of control or BLTP1 KO cells from three independent assays. The mean H2DCFDA fluorescence intensity was normalized to the control. More than 10000 cells for each condition were analyzed. Two-tailed unpaired Student’s t test. Mean ± SD. Scale bar, 10 μm in (D, E).

**Fig. S5.**
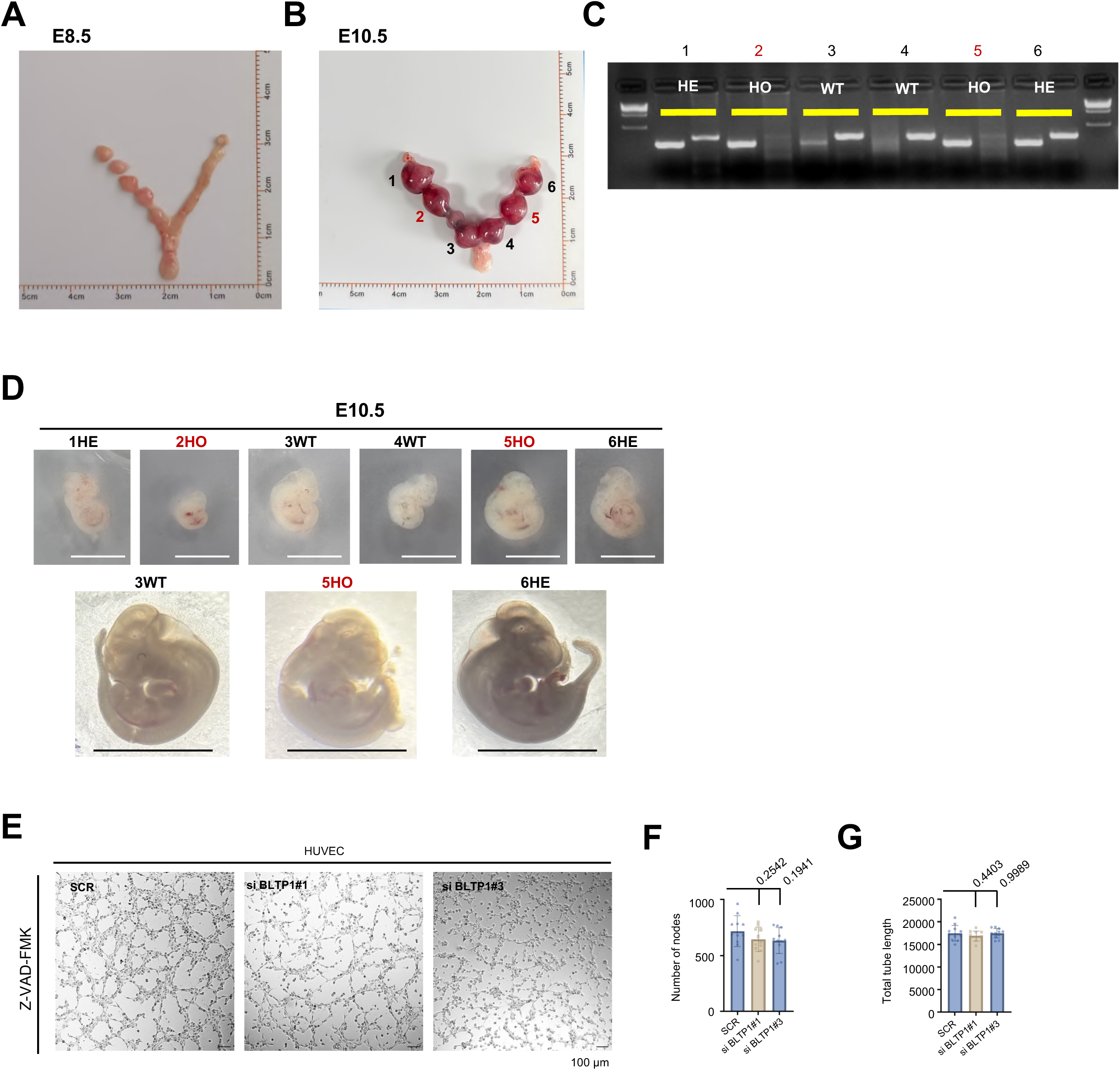
LPD-3 KO mediated by CRISPR-Cas9 caused developmental defects. (A) CRISPR-Cas9-mediated KO of LPD-3 in animals [LPD-3(-)]. Two sgRNAs are used with the underlined letters, indicating the PAMs (CGG for sgRNA1 and TGG for sgRNA2) for spCas9. (B) Genotyping confirming single animals for WT and LPD-3(-) candidates. (C) Representative fluorescent images of WT or LPD-3 (-) animals in the pgl-1 (ax3122) strain at stages of L4+24h, L4+48h or L4+56h. (D) Normalized length of LPD-3 (-) animals as in (C). More than 10 animals for each condition were analyzed. Two-tailed unpaired Student’s t test. Mean ± SD. (E) Quantification of degree of gonadal deformation of LPD-3 (-) animals at multiple developmental stages from three independent experiments. More than 16 animals for each condition were analyzed. (F) Representative bright-field images of control or LPD-3(-) animals in the pgl-1(ax3122) strain. (G) Quantification of numbers of progeny of WT (n=9) or LPD-3 (-) (n=9) animals from three independent experiments. Two-tailed unpaired Student’s t test. Mean ± SD. Scale bar, 100 μm.

**Fig. S6.**
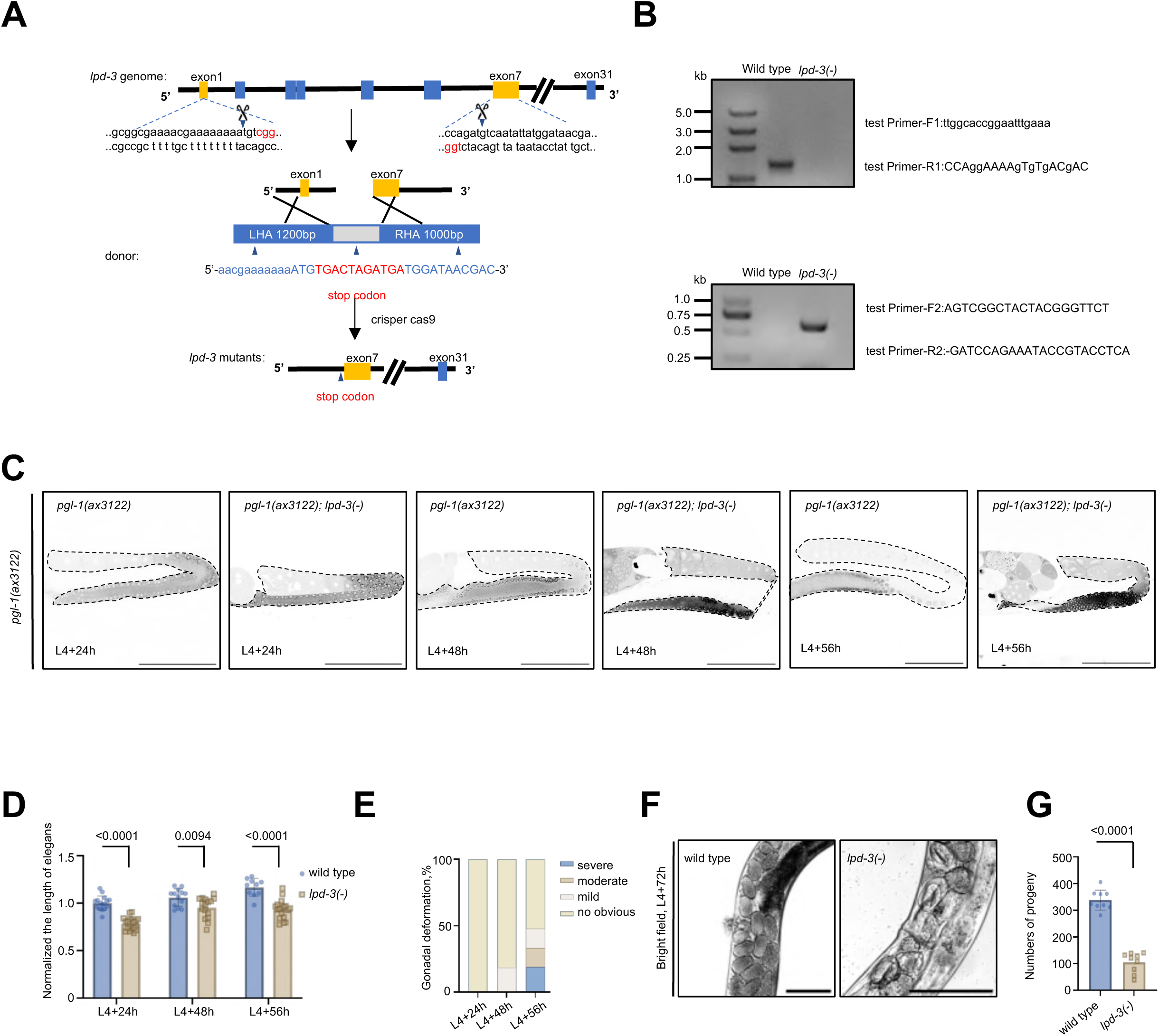
BLTP1-/- mice did not exhibits severe defect in embryo at ED 8.5 or 10.5. (A, B) The morphological features of uteri at 8.5 (A) or 10.5 (B) days post-fertilization. (C) DNA gel showing genotypes of WT, HE, and HO BLTP1 KO mice embryos. (D) Representative images of six embryos as in (B) with HO embryos (2-HO and 5-HO) being labeled in red. (E) Representative images of tube formation assays of control or BLTP1 depleted HUVEC cells in presence of apoptosis inhibitor Z-VAD-FMK (20 μM). (F, G) Node number (F) or tube length (G) in cells as in (E). 11 ROIs from three independent experiments were quantified. Ordinary one-way ANOVA with Tukey’s multiple comparisons test. Mean ± SD. Scale bar, 0.5 cm in (D) and 100 μm in (E).

**Fig. S7.**
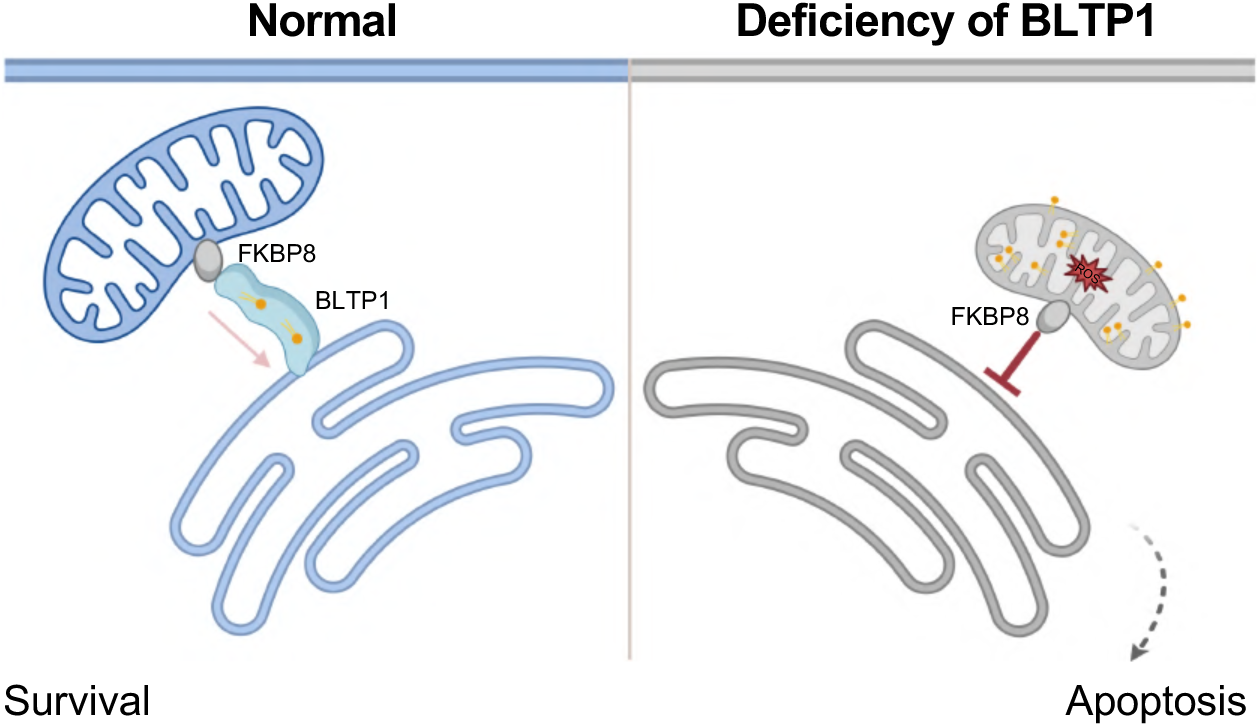
The working model of BLTP1 functions at ER-mitochondria contacts. BLTP1 is an ER integral protein that is essential for life. FKBP8 recruits BLTP1 to ER-mitochondrial contacts. At these sites, BLTP1 transports phospholipids (e.g., PG and PA) out of mitochondria and maintains mitochondrial lipid homeostasis by counterbalancing the influx of lipids required for CL synthesis. Its deficiency triggers a cascade of mitochondrial lipid overload, oxidative stress, dysfunction, and apoptosis.

